# Prolonged tonic pain in healthy humans enhances functional connectivity of descending pain modulation networks involving the amygdala, periaqueductal gray and parabrachial nucleus to cortical sensory-discriminative areas

**DOI:** 10.1101/2021.08.31.458440

**Authors:** Timothy J. Meeker, Anne-Christine Schmid, Michael L. Keaser, Shariq A. Khan, Rao P. Gullapalli, Susan G. Dorsey, Joel D. Greenspan, David A. Seminowicz

## Abstract

**Introduction:** Resting state functional connectivity (FC) is widely used to assess functional brain alterations in patients with chronic pain. However, reports of FC changes accompanying tonic pain in pain-free persons is rare. A brain network disrupted during chronic pain is a network we term the Descending Pain Modulatory Network (DPMN). Here, we evaluate the effect of tonic pain on FC of this network: anterior cingulate cortex (ACC), amygdala (AMYG), periaqueductal gray (PAG), and parabrachial nuclei (PBN).

**Methods:** In 50 pain-free participants (30F), we induced tonic pain using a capsaicin-heat pain model. We used functional MRI to measure resting BOLD signal during pain-free rest where participants experienced warmth and tonic pain where participants experienced the same temperature thermode combined with capsaicin. We evaluated FC from ACC, AMYG, PAG, and PBN with correlation of self-report pain intensity with FC during both states. We hypothesized tonic pain would disrupt FC dyads within the DPMN. We used partial correlation to determine FC correlated with pain intensity and BOLD signal.

**Results:** Of hypothesized FC dyads, PAG and subgenual ACC was weakly disrupted during tonic pain (F=3.34; p=0.074; pain-free>pain d=0.25). sgACC-PAG FC became positively related to pain intensity (R=0.38; t=2.81; p=0.007). Right PBN-PAG FC during pain-free rest positively correlated with subsequently experienced pain (R=0.44; t=3.43; p=0.001). During tonic pain, FC of this connection was abolished (paired t=-3.17; p=0.0026). During pain-free rest, FC between left AMYG and right superior parietal lobule and caudate nucleus were positively correlated with subsequent pain. During tonic pain, FC between left AMYG and right inferior temporal and superior frontal gyri negatively correlated with pain. Subsequent pain positively correlated with right AMYG FC and right claustrum; left and right primary visual cortex; right middle temporal gyrus and right temporo-occipitoparietal junction. Finally, subsequent pain positively correlated with PAG FC and left cerebellum, left dorsolateral prefrontal, right posterior cingulate cortex and paracentral lobule, inferior parietal lobule, medial precuneus and PBN.

**Conclusion:** We demonstrate 1) tonic pain weakly disrupts of sgACC-PAG FC; 2) sgACC-PAG tonic pain FC positively correlates with pain; 3) right PBN-PAG FC predicts subsequent pain and is abolished during tonic pain. Finally, we reveal PAG- and right AMYG-anchored networks which predict intensity of tonic pain. Our findings suggest specific connectivity patterns within the DPMN at rest predict experienced pain and are modulated by tonic pain. These nodes and their functional modulation may reveal new therapeutic targets for neuromodulation and biomarkers to guide interventions.

**Highlights:**

- Parabrachial-periaqueductal gray (PAG) functional connectivity (FC) predicts pain
- Subgenual anterior cingulate cortex-PAG FC correlates with pain during tonic pain
- PAG- and amydalocortical networks at rest predict tonic pain intensity
- Resting FC of PAG supports cortical targets of neuromodulation to control pain

## Introduction

Functional connectivity (FC) has emerged over the past two decades as a technique to investigate the functional anatomy of human brain networks and the effects of psychological, pathological and perceptual manipulations, therapeutic treatments, and disease states on brain function (Baliki et al., 2012; Biswal et al., 1995; Crowther et al., 2015; Khalili-Mahani et al., 2017; Raichle et al., 2001). A network of brain regions, which we term the descending pain modulatory network (DPMN), has been well-studied during simultaneous experience of phasic painful heat stimuli, concurrent painful heat and distracting stimuli, during mind-wandering, placebo analgesia, and altered pain states such as brush allodynia (Becerra et al., 2006; Bingel et al., 2006; Eippert et al., 2009; Kucyi et al., 2013; Linnman et al., 2012; Moulton et al., 2007; Petrovic et al., 2002; Valet et al., 2004). We consider a cortical or subcortical brain region to be part of the DPMN if it is differentially activated during pain modulation and has relatively high concentrations of µ-opioid receptors (Henriksen and Willoch, 2008). Demonstrating the potential clinical importance of the DPMN, enhanced FC within the DPMN during placebo analgesia and motor cortex stimulation is positively related to magnitude of analgesia experienced (Bingel et al., 2006; Eippert et al., 2009; Garcia-Larrea and Peyron, 2007; Meeker et al., 2019); additionally, there is a reduction of FC of the DPMN in chronic pain patients compared to pain-free controls (Linnman et al., 2012; Yu et al., 2014). In healthy individuals, coactivation of the PAG and ACC during the experience of phasic pain coupled with analgesic cognitive manipulations, such as placebo, results in enhanced FC between these regions accompanied by a reduction in perceived pain intensity (Bingel et al., 2006; Eippert et al., 2009; Fairhurst et al., 2007; Kucyi et al., 2013; Sprenger et al., 2011; Valet et al., 2004). Several regions in the ACC surrounding the genu of the corpus callosum have been implicated in pain modulation, therefore we interrogate three separate ROIs in the region of the ACC including pregenual (pACC), anterior subgenual ACC (sgACC) and supragenual ACC (spACC) (Bingel et al., 2006; deCharms et al., 2005; Seymour et al., 2005; Valet et al., 2004; Wiech et al., 2005).

While several studies have evaluated the network’s FC during phasic painful stimuli there is a scarcity of studies exploring FC during prolonged tonic pain (Ayoub et al., 2021; Bingel et al., 2006; Kucyi et al., 2013; Seminowicz and Davis, 2007; Valet et al., 2004). Tonic pain states, such as those encountered in chronic pain syndromes, display unique perceptual dynamics and modeling prolonged tonic pain in healthy participants is a critical intermediate step in understanding the neurophysiology of chronic pain disorders (Baliki et al., 2006; Foss et al., 2006).

To probe the functional modulation of relationships within the DPMN during a prolonged tonic painful stimulus in a preclinical human pain model, we acquired resting state fMRI scans in pain-free participants before and after exposing them to a potent topical capsaicin-heat pain (C-HP) model (Anderson et al., 2002; Meeker et al., 2019), thereby capturing pain-free and prolonged tonic pain resting states within a single imaging session.

In this report, we predicted that prolonged tonic pain would disrupt the coupling between the ACC and PAG since disruption in FC between ACC and PAG occurs in chronic pain disorders and the ACC displays reductions in local FC in chronic pain (Ke et al., 2015; Khalili-Mahani et al., 2017; Li et al., 2016; Liu et al., 2012; Wei et al., 2016; Wu et al., 2016). We further predicted enhanced FC between the AMYG and PBN during tonic pain compared to the pain-free state given the amygdaloparabrachial transmission pathway and its role in pain response and modulation as evidenced in rodent studies (Chen and Heinricher, 2019; Kissiwaa and Bagley, 2018; Raver et al., 2020; Roeder et al., 2016; Uddin et al., 2018). Finally, we predicted modulation of a functional connection between the PAG and parabrachial nucleus, given primate tractography and rodent neurophysiology implicating this pathway in pain modulation (Chen and Heinricher, 2019; Mantyh, 1982a, 1983; Roeder et al., 2016). Evidence supporting these predictions would suggest that in chronic pain, ongoing/spontaneous pain modulates resting state FC. Importantly, this could contribute to FC measures of disease severity and classification independently of disease progression. Objective metrics derived from FC measures related to pain intensity developed in these models could be used to predict populations susceptible to chronic pain to allow additional preventive treatments in these susceptible patients. Finally, modulation of FC between pain modulatory areas may be tracked during clinical trials to reveal intervention mechanisms and provide objective measures to track treatment progress (Borsook et al., 2012; Meeker et al., 2019).

## Methods

### Overview

We report results from a total of 50 participants enrolled in one of two studies conducted at University of Maryland Baltimore (UMB) from October 2011 until December 2015. Study 1 set out to establish the effects of a prolonged tonic pain stimulus lasting several minutes on the functional organization of the human brain. In the first experiment, we conducted experiments provoking pain in 18 healthy participants (10 M; 2 left-handed) aged 23 to 61 (median=30.5) by applying 10% capsaicin cream under a warm thermode on their left leg, which we term the capsaicin-heat pain (C-HP) model to distinguish it from previous models using lower concentration capsaicin creams (Anderson et al., 2002; Cavallone et al., 2013; Meeker et al., 2019; Petersen and Rowbotham, 1999). We conducted a screening session to eliminate participants who did not develop sufficient heat allodynia during the C-HP (Liu et al., 1998). Eligible participants then took part in an MRI session which was separated by ≥13 days (median=39.5 (range=13 to 88)) from the screening session. Using the C-HP model we maintained a mild to moderate pain intensity with a 39°C (n=11), 40 °C (6) or 41 °C (1) thermode. The temperature selected for each participant was based on that individual’s thermal heat pain sensitivity evaluated at screening just prior to the C-HP exposure.

In study 2, 40 participants (17 M; median age: 24; range 20-39) underwent an MRI in which we employed the C-HP model with a 38°C (n=4), 39 °C (2), 40 °C (6), 41 °C (8), or 42 °C (20) thermode. From this group of participants, 3 males were excluded from the current report because they reported pain ratings of 0 out of 100 during the last 2 minutes of exposure to the C-HP model (C-HP temperatures for these participants were all 42 °C). A further 4 males and 1 female were excluded due to having greater than 0.5 mm motion framewise displacement in at least 10% of the functional MR time series (1 at 40 °C, 1 at 41 °C, 2 at 42 °C). Resting state fMRI results from study 1 and 2 are pooled together and use the same MRI sequence and protocol, excepting that scans were acquired at different resolutions (1.8 x 1.8 x 4 mm^3^ versus 3 mm^3^ isotropic) and different head coils (12-versus 32-channel). All participants provided written informed consent, and all procedures were approved by the UMB Institutional Review Board for the Protection of Human Subjects.

### Eligibility Criteria

In study 1 exclusion criteria were: pregnancy; history of brain injury with any period of unconsciousness; illicit, or prescription opioid, drug use; current pain or history of chronic pain; history of cardiac, renal, hepatic, or pulmonary function disorders; history of cancer; ambidextrous (Oldfield, 1971); hospitalized for a psychiatric disorder within last 12 months; pain intensity rating less than 21 on a 0-100 NRS while exposed to the C-HP model. Illicit drug use was determined with a urine drug screen for marijuana, cocaine methamphetamine, amphetamines, ecstasy, heroin, phencyclidine, benzodiazepines, methadone, barbiturates, tricyclic antidepressants or oxycodone (First Check™).

In study 2, in addition to eligibility criteria for study 1, we excluded left-handed participants, any individual with any diagnosis of psychological or neurological disorder or participants taking any psychoactive medications (by self-report). However, in study 2 we did not exclude any individual based on sensitivity to the C-HP model.

### Psychophysics and Psychological Questionnaires

During the initial session of each study, we measured participants’ warmth detection thresholds (WDTs) and heat pain thresholds (HPTs) with a Medoc stimulator (Pathway; Medoc; Ramat Yishai, Israel) using the method of limits (Greenspan, 2013). We placed the 3×3 cm contact area stimulator on the lower left foreleg at a baseline temperature of 32 °C. A program increased the temperature at a ramp of 0.5 °C/s until the participant pressed a mouse button. We instructed the participant to press the button when they “felt a change in temperature” for WDTs or when the warmth “becomes painful” for HPTs. At a single site, we measured four trials for WDTs and HPTs. We took the average of the last three threshold determinations for each participant. In study 2, the protocol for HPTs started with a baseline of 30 °C, to accommodate sensitivity changes after capsaicin exposure.

### Capsaicin-Heat Pain Model

In order to produce a safe, sustained painful experience, we treated participant’s lower left foreleg with one gram of 10% capsaicin cream under a Tegaderm™ bandage (Anderson et al., 2002; Meeker et al., 2019). To control the exposure area, we applied the cream within a 2.5 cm^2^ square cut into a Tegaderm™ bandage. After 12 minutes of exposure – long enough for the capsaicin cream to reach saturating concentrations at the intraepidermal nerve fiber endings – we placed the thermode over the topmost bandage at the designated temperature (Green and Flammer, 1988). During study 2 the incubation period was increased to 15 minutes and the thermode was placed on the participant’s leg during incubation, held at 32 °C. Target temperatures used were tailored for each participant between the pre-capsaicin WDT and HPT. Participants rated pain intensity on a numerical rating scale (NRS) with verbal anchors on one side, and numbers ranging from 0-100 in increments of 10 (Greenspan et al., 2003). In study 1, participants provided pain intensity ratings every 30 seconds for 10 minutes after application of the thermode. Participants reporting average NRS pain 30 out of 100, and tolerating the C-HP model, were eligible for MRI sessions. During study 2 participants provided pain intensity ratings every minute for 35 minutes during the entire capsaicin exposure. At the end of the exposure period, we removed the bandages and capsaicin with an isopropanol swab. This C-HP procedure does not cause tissue damage (Moritz and Henriques, 1947).

### MRI Procedures

We recorded fMRI in a 3-T Tim Trio scanner (Siemens Medical Solutions, Malvern, PA) using a 12-channel (study 1) or 32-channel (study 2) head coil with parallel imaging capability. For resting state scans during study 1, we used a gradient echo single-shot echo-planar-imaging (EPI) sequence with 30 ms echo time (TE), 90° flip angle and 2500 ms repetition time (TR) providing T2*-weighted volumes in 36 interleaved, 4 mm slices (no gap) with an in-plane resolution of 1.8 mm^2^. For study 2, we used identical parameters except that we collected 44 interleaved, 3 mm slices (no gap) with an in-plane resolution of 3.0 mm^2^. During both resting state scans participants fixated on a crosshair for 8 min 12.5 s providing 194 functional volumes. For anatomical reference, we acquired a 3-dimensional T1 magnetization-prepared rapid gradient echo (MPRAGE) volumetric scan with 2.9 ms TE, 2300 ms TR, 900 ms inversion time (TI), flip angle 9°, 144 slices, axial slice thickness 1.0 mm and 0.9 mm2 in-plane resolution over a 23-cm field of view for 13 of 14 participants belonging to study 1. The remaining one participant from study 1 and all participants from study 2 received a modified MPRAGE acquisition which facilitated isotropic resolution and provided extended coverage of the brain: 2.9 ms TE, 2300 ms TR, 900 ms TI, flip angle 9°, 176 slices, sagittal slice thickness 1.0 mm and 1.0 mm^2^ in-plane resolution over a 25.6-cm field of view. Since structural scans were used for anatomical reference and display only, this difference in anatomical acquisition did not influence the results.

### fMRI Session Protocol

During study 1 and 2 MRI sessions, we evaluated the participants’ WDTs and HPTs in the MRI environment. During the resting state scan participants were told: “Please stare at the plus sign, do not move and do not fall asleep. You may let your mind wander.” Then, we increased the thermode temperature to the predetermined target, while the participant fixated on the cross hair for the duration of the scan. After the scan, participants remained in the scanner with the thermode in place and rated their pain intensity every 30 seconds for two minutes on a 0-100 NRS.

### Statistical Methods

Effects of time on sensory detection thresholds (WDTs and HPTs), after exposure to the C-HP model, were evaluated using a linear mixed model with time as a fixed factor and participant as a random factor. In different models, either baseline HPT or baseline WDT were using as a control to compare to HPTs taken at 25, 50 and 75 minutes after capsaicin removal. Multiple comparisons were corrected using Tukey’s HSD. The R package ‘anova’ was used to derive F-stats for the overall model. We conducted this set of statistical tests using R 3.4.1.

### Resting state fMRI data analysis

All preprocessing of resting state fMRI scans used the afni_proc.py python script for Analysis for Functional NeuroImaging (AFNI) version 27 Jun 2019. The first three volumes were automatically removed from the functional scan series by the MRI scanner to allow for signal equilibration. We used 3dToutcount to determine the minimum outlier EPI volume for later EPI volume registration and alignment. Outliers were defined in relation to the median absolute deviation of the signal time course (see afni.nimh.nih.gov/pub/dist/doc/program_help/3dToutcount.html for outlier definition). Each functional time series was detrended and spikes quashed with 3dDespike. Then, each volume was slice-time corrected and aligned to the first slice collected during the TR. Before aligning the anatomical scan to the functional scan, the skull was removed from each individual Freesurfer processed anatomy using 3dSkullStrip. We used 3dAllineate via the align_epi_anat.py python script to align the anatomy to the minimum outlier functional EPI volume using the local Pearson correlation signed cost functional while allowing different cost functionals for initial alignment if, for example, alignment fails (e.g., lpc+ZZ). After alignment, the anatomical volume was warped to MNI atlas space and normalized to the MNI152, version 2009, skull-stripped brain using @auto_tlrc. Each EPI volume was then registered to the base, minimum outlier, EPI volume. Then, the registered EPI volumes were aligned to the template-aligned structural volume using non-linear warping. Following this final functional alignment, a full alignment matrix was estimated and applied to the anatomy follower dataset from Freesurfer including default Freesurfer parcellations (e.g., aaseg) and ventricle and white matter segments, which were first eroded by 1 mm^3^. Since the DPMN, as we have defined it, involves several midbrain and brainstem regions, each 4D EPI dataset was then blurred with a 4 mm FWHM Gaussian spatial filter. To provide a normalized across participant standard interpretation of signal fluctuations and prevent any participant from outweighing any other participant, we scaled the 4D BOLD signal to a normalized value of a mean of 100 and range from 0-200.

In the participant-level regression model regressors of no interest included a binary regressor excluding volumes with motion exceeding 0.5 mm in framewise displacement, the principal component of signal extracted from the individual eroded Freesurfer ventricle masks, demeaned motion parameters (motion in x, y and z planes and rotation about the x, y and z axes), and their first order derivatives. There were 72 out of 9700 (0.74%) volumes censored during the pain-free resting state scan and 118 out of 9700 (1.22%) volumes censored during the pain state scan. We did not regress out any signal or signal-derivative from white matter since this may have removed variance from gray matter brainstem regions. An LMM of average motion as framewise displacement for each participant with factors of sex and state found no significant effect of state (F=1.99; p=0.165), sex (F=0.75; p=0.389), or state-by-sex interaction (F=2.32; p=0.134). Pain intensity during the pain state was not related to average framewise displacement during the pain-free (R=-0.031; t=-0.22; p=0.83) or pain state scan (R=0.031; t=0.22; p=0.83). For each participant, the regressor of interest was the seed time course for that specific seed and session. We have divided our report into cortical, subneocortical and basal ganglia regions of interest. Here we report the results of the subcortical investigation including the periaqueductal gray (PAG), left and right AMYG and left and right parabrachial regions. We exclude the dorsal raphe and rostral ventromedial medulla as seeds as specific fMRI acquisition methods are needed to optimize acquisition of signal from the medulla (Beissner et al., 2014; Stroman et al., 2018). The seed regions of interest were left and right AMYG as defined by Freesurfer parcellation, three bilateral 5 mm radius ROIs in the anterior cingulate cortex corresponding to supragenual (MNI: ±6, (median=35; min=33; max=37), (median=12; min=10; max=14)), pregenual (MNI: ±6, (median=38; min=37; max=40), (median=1; min=-2; max=3)) and subgenual (MNI: ±5, (median=31; min=29; max=37), (median=-9; min=-11; max=-7)) ACC, and 3 mm radius spheres for left and right parabrachial nucleus (PBN) in (MNI: x=±10, y=-35, z=-30), and a participant-specific anatomically drawn seed for the PAG (PAG; group average center of mass: MNI: (0, - 29, -10)) (Fairhurst et al., 2007; Linnman et al., 2012; Morey et al., 2009; Sprenger et al., 2011). The PAG seed was drawn to encompass all apparent gray matter surrounding the cerebral aqueduct on T1 MPRAGE images after alignment into atlas space. ACC seeds were placed on individual participant anatomies and tests completed to ensure they did not overlap and were placed within gray matter. Seeds that were drawn or placed anatomically by TJM (L- and RPBN, spACC, pACC, sgACC and PAG) were each reviewed for placement by both TJM and JDG on individually aligned anatomical images. Since few reports of network brain connectivity in humans including the whole brain have reported FC in the context of tonic pain in otherwise healthy participants, we include exploratory analysis of each seed region including contrasts and with FC covariation with pain intensity experienced during tonic pain (Ayoub et al., 2021; Pritchard et al., 2000; Stroman et al., 2018).

Seed-to-seed FC analysis was performed on the R-to-Z-transformed FC of the following dyads: spACC-PAG, pACC-PAG, sgACC-PAG, spACC-left amygdala (LAMYG), spACC-right amygdala (RAMYG), pACC-LAMYG, pACC-RAMYG, sgACC-LAMYG, sgACC-RAMYG, sgACC-LPBN, sgACC-RPBN, LPBN-LAMYG, RPBN-RAMYG, LPBN-PAG, RPBN-PAG, sgACC-LPBN, and sgACC-RPBN. Each LMM included participant as a random effects factor, and state (tonic pain vs. pain-free) and sex (M vs. F) as fixed effects factor and the interaction of state and sex. Specifically for these dyads, supported by extensive tract-tracing evidence from animal models and prior FC findings in either healthy and chronic pain patients or functional coupling reactive to nociceptive stimuli in animal models, we accepted one-sided directional significance tests (p<0.1) based on the hypothesis that these connections would be disrupted during the pain state when compared to the non-pain state. We point out when our directional assumptions are violated and the p-values reported would be identical under either uni- or bi-directional hypothesis tests. It should be noted the guidelines ‘significance’ of p-values, are arbitrary and participant to test assumptions, sample size, context, and practical effect interpretation(Fisher, 1948; Wasserstein and Lazar, 2016). Degrees of freedom for F-tests and posthoc analyses were corrected for using the Satterthwaite correction (Luke, 2017). Additionally, we calculated Pearson correlation between normalized, mean-centered pain intensity ratings during the tonic pain state with each FC dyad for each state. We report both uncorrected p-values and if they are significant after Bonferroni correction for the 30 dyad-state pairings and 15 contrasts between states for each dyad, as a subtraction of t-stats of the R-scores calculated on each dyad. This analysis was completed with R version 3.6.3.

### Group Level whole-brain fMRI Analysis

For group level analysis, we used AFNI’s linear mixed-effects modeling program 3dLME (Chen et al., 2013). A two-factor model focused on the change in state between the pain-free and prolonged tonic pain resting states with the C-HP model inducing heat allodynia (factor levels: pain-free and tonic pain) to arrive at tonic pain - pain-free contrast maps and mean seed-driven functional connectivity network (FCN) maps. For pain intensity covariation with seed-driven, FC we used AFNI’s 3dttest++. All analyses were restricted within a group gray matter probability mask which excluded ventricles and cortical white matter regions. First each individual gray matter mask was created by subtracting the individual binary Freesurfer ventricle and cortical white matter masks from the group union functional mask. Then the group binary gray matter probability mask was created by including voxels where at least 30 of 50 individual participants had gray matter. We implemented a minimal voxel-wise p-value threshold of 0.005 for covariate analyses and 0.001 for contrast analyses. For FCN maps of brainstem and midbrain seeds we cluster-extent corrected for R-score and p-value, thresholding at R=0.10 and p=0.0001. For FCN maps of amygdala seeds we cluster-extent corrected for R-score and p-value, thresholding at R=0.20 and p=0.0001. To correct for multiple comparisons, we estimated the spatial autocorrelation function of the residual noise of the BOLD signal within our analysis mask using 3dFWHMx and used the resulting parameters with 3dClustSim to calculate cluster extent criteria (CEC) for both 3dLME and 3dttest++ resultant maps.

For voxel tables detailing covariate results, we implemented an initial voxel-wise threshold of 0.005, while contrast results voxel tables had an initial voxel-wise threshold of 0.001. To elucidate the coordinates of local maxima within large clusters, we iteratively reevaluated statistical maps after reducing the p-value threshold of each map by a factor of 10 (e.g., 0.001-0.0001). This process was completed by hand after generating all possible iterative voxel tables using 3dclust. For group FCN maps voxel tables, we implemented an initial minimum cluster-extent corrected for R-score and p-value, thresholding at R=0.10 (R=0.20 for AMYG seeds) and p=0.0001. We followed a similar iterative process for FCN voxel tables as for contrast and covariate voxel tables but increased the R-score in 0.05 steps.

Following the pain intensity covariate analysis, we calculated the partial correlation matrix of the brain regions which correlated with pain intensity. This allowed us to test for significant correlations among the BOLD signal among these brain regions while 1) controlling for pain intensity correlation and 2) investigating the interdependence among brain regions and the seed ROI controlling for network interdependencies. Before this analysis we instituted an R-to-Z transform on the pain intensity correlations with FC metrics. We completed this analysis using R package ‘ppcor’ (Kim, 2015). Furthermore, we used R (version 3.6.3) to create correlation plots of pain intensity to the BOLD FC between the seed ROI and significant regions discovered in the whole-brain analysis.

## Results

### Psychophysical and Perceptual Response to Capsaicin-Heat Pain (C-HP) Model During Screening Session

All participants included in this analysis reported hot-burning pain in response to the C-HP model. For participants in study 2 (n=32), the time course for pain intensity ratings during the screening session shows an ever-increasing trend during the C-HP exposure, which responded positively to 0.5° C step increases at each arrow (Fig. 1A). The thermode temperature for the MRI scan was tailored for each participant, and once increased to the target temperature, remained stable during the MRI sessions. Further, in study 2 (n=32), exposure to the C-HP model induced profound heat allodynia which lasted at least 75 minutes after capsaicin removal (F=11.28; p<0.001) (Fig 1B). HPTs after capsaicin removal were all significantly lower than pre-capsaicin exposure HPTs (t≤ -7.8; p≤10^-8^) and the first HPTs after C-HP removal were lower than baseline WDTs (t=-4.5; p=8.4×10^-5^), demonstrating profound hypersensitivity to heat after C-HP exposure. Using SFMPQ-2 descriptors, most participants described the pain as throbbing, aching, heavy, tender, shooting, stabbing, sharp, piercing, sensitive to touch, hot-burning and tingling (Fig 1C). After removing the descriptor ‘hot-burning’ from the analysis, we found a significant effect of pain descriptor class (F=9.17; p<0.0001) and no effect of sex on participantive ratings on the SFMPQ-2 (F=0.18; p=0.67). Participants rated both intermittent and continuous pain descriptors higher than either neuropathic or affective pain descriptors (Continuous>Affective: t=4.0; p<0.001; Intermittent>Affective: t=4.0; p<0.001; Continuous>Neuropathic: t=4.1; p<0.001; Intermittent>Neuropathic: t=4.1; p<0.001). There was no difference in ratings between continuous and intermittent descriptors, and neuropathic and affective descriptors (t<0.13; p>0.98). During the MRI experiment, during the two minutes immediately after the corresponding tonic pain resting state scan average pain intensity of the group was rated as 33 (SD=21) on a 0-100 point scale with a range of 1-75.

**Figure 1.**
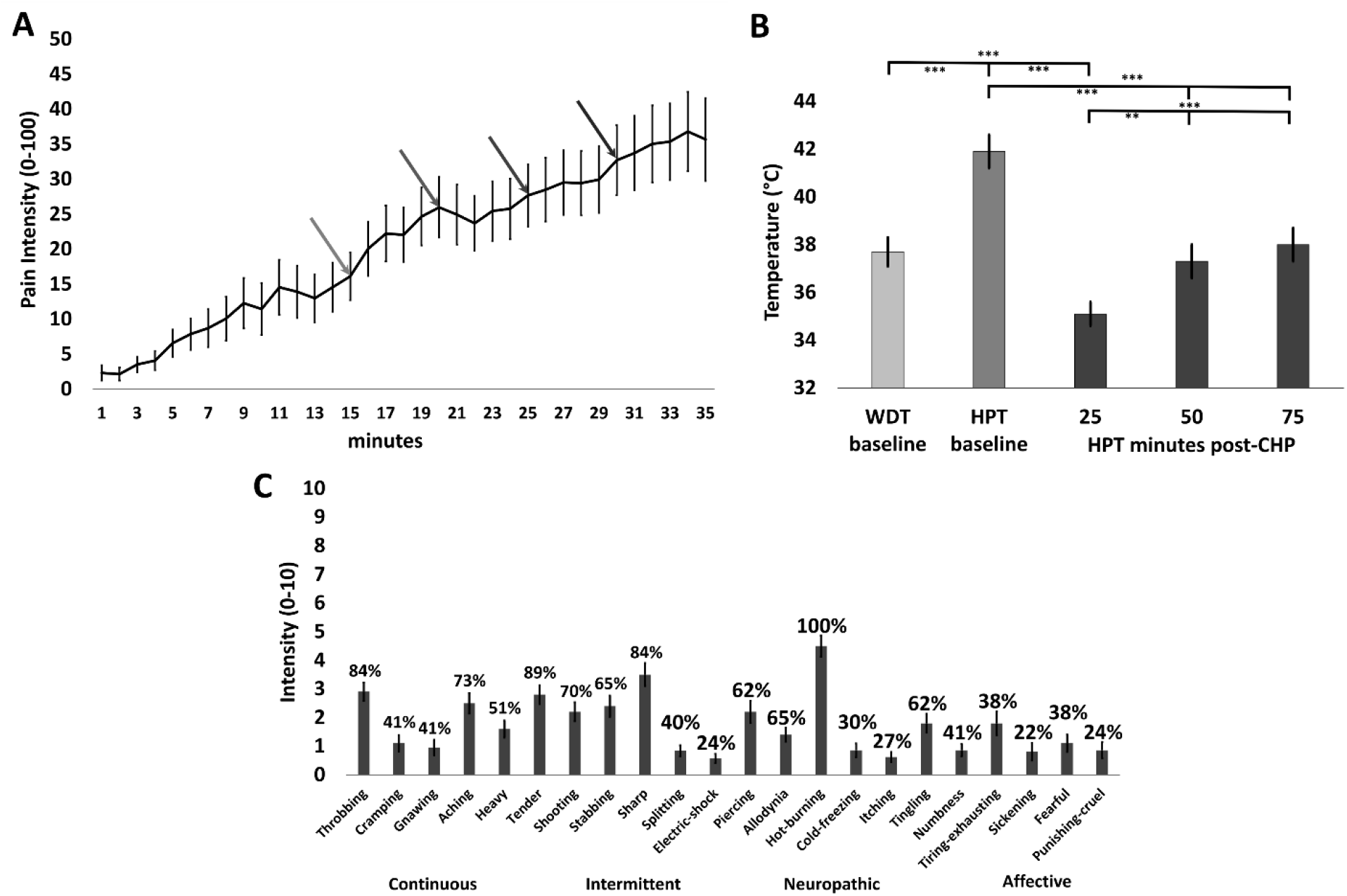
**A)** Ratings of prolonged tonic pain intensity during C-HP, the first arrow is when the thermode increased to the target temperature and the temperature increased 0.5°C at each subsequent arrow (n=32). **B)** Warmth detection thresholds (WDT) and heat pain thresholds (HPT) taken before and after exposure to the C-HP model (n=32); ** - p=0.0002; *** - p < 0.0001. **C)** Mean intensity of pain descriptors from the SFMPQ-2, where percentages above each bar represent percentage of subjects endorsing pain descriptor with any non-zero rating. All error bars are SEM.

### Resting State Functional Connectivity: Effects of tonic pain on specific DPMN dyads

For the LMM analysis including sex, state and sex-state interaction effects, connectivity of the three ACC seeds to the PAG, sex was a significant factor for the pACC-PAG connection (F=4.84; p=0.030; M>F Cohen’s d=0.59) and state was a significant one-sided factor for the sgACC-PAG connection, where the pain state disrupted this connection (F=3.34; p=0.074; pain-free>pain d=0.25) (Fig. 2A). State was not significant for either the pACC-PAG (F=0.66; p=0.42) or spACC-PAG (F=0.16; p=0.69) connections. Sex was not a significant factor for either the sgACC-PAG (F=2.23; p=0.14) or spACC-PAG (F=0.036; p=0.85) connections. The interaction between sex and state was not a significant for the pACC-PAG (F=0.93; p=0.34), spACC-PAG (F=0.90; p=0.35) or sgACC-PAG (F=0.54; p=0.47) connections. For the two functional connections tested between the bilateral sgACC and left and right PBN, neither sex, state nor the sex by state interaction was significant for the sgACC-LPBN (sex: F=0.000; p=1.00, state: F=0.67; p=0.42, sex-by-state: F=0.072; p=0.79) or sgACC-RPBN (sex: F=0.005; p=0.95, state: F=0.008; p=0.93, sex-by-state: F=0.49; p=0.49).

**Figure 2.**
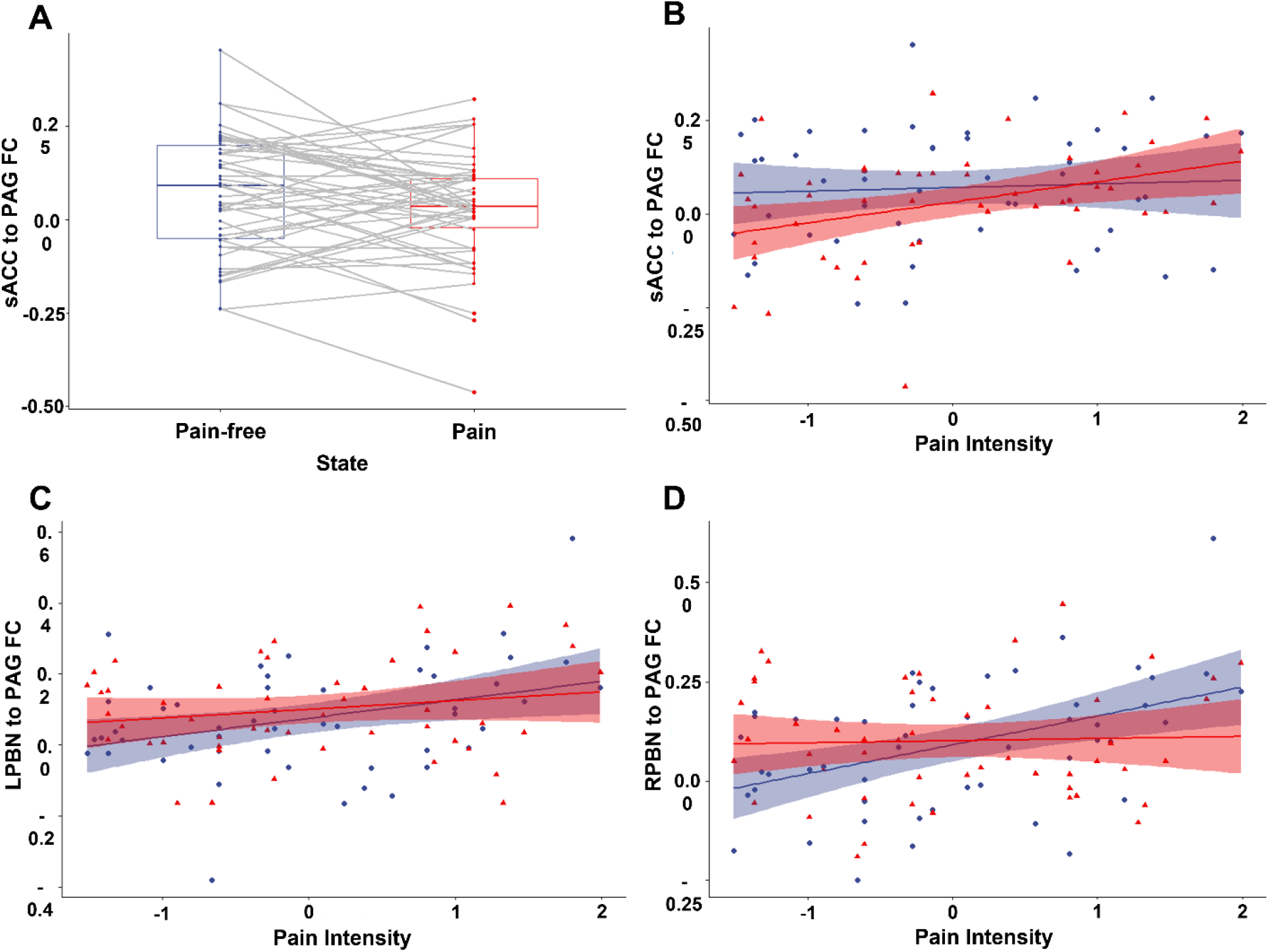
**A)** Change in functional connectivity between subgenual anterior cingulate cortex and periaqueductal gray from pain-free to tonic pain state. **B)** Correlation between normalized pain intensity and functional connectivity between subgenual anterior cingulate cortex and periaqueductal gray during the pain-free (blue circles) and tonic pain states (red triangles). **C)** Correlation between normalized pain intensity and functional connectivity between left parabrachial nucleus and periaqueductal gray during the pain-free (blue circles) and tonic pain states (red triangles). **D)** Correlation between normalized pain intensity and functional connectivity between right parabrachial nucleus and periaqueductal gray during the pain-free (blue circles) and tonic pain states (red triangles). Shaded areas correspond to 95 percent confidence bounds.

For the four connections from the left and right PBN, neither sex, state nor the sex by state interaction was significant for the LPBN-PAG (sex: F=0.006; p=0.95, state: F=2.52; p=0.12, sex-by-state: F=1.60; p=0.21), LPBN-LAMYG (sex: F=2.04; p=0.16, state: F=0.79; p=0.38, sex-by-state: F=0.002; p=0.96), or RPBN-RAMYG (sex: F=0.68; p=0.41, state: F=1.25; p=0.27, sex-by-state: F=0.33; p=0.57) connections. However, while sex (F=1.35; p=0.25) and state (F=2.99; p=0.09) were not significant effects on the FC of the RPBN-amygdala connection, the sex-by-state interaction was significant (F=4.69; p=0.035). This resulted from FC in this connection decreasing during the tonic pain state compared to the pain-free state in females (n=30; d=0.30) and increasing during the tonic pain state compared to the pain-free state in males (n=20; d=0.30). However, posthoc comparisons were not significant after multivariate t correction (t<1.73; p>0.31). For the six functional connections tested between the bilateral ACC and left and RAMYG, neither sex, state nor the sex by state interaction was significant for the ACC-LAMYG (sex: F<2.22; p>0.14, state: F<0.89; p>0.35, sex-by-state: F<0.25; p>0.62) or ACC-RAMYG (sex: F<1.36; p>0.25, state: F<3.26; p>0.077, sex-by-state: F<1.58; p>0.21). While the state effect on the spACC-RAMYG functional connection reached one-sided significance, the a priori directional nature for the effect of pain was opposite to our predictions, therefore it was not considered significant (Cohen’s d=0.20; pain state > pain-free state FC).

We tested for correlations of pain intensity during the tonic pain experience for with each FC dyad during both the pain and pain-free states (Table 1). Before induction of the tonic pain model correlation between pain intensity and FC between the sgACC and PAG was not significant, however, during the tonic pain experience correlation between pain intensity and FC between the sgACC and PAG became positive (R=0.38; t=2.81; p=0.007) (Fig. 2B), surviving BF correction (over 30 dyads). Furthermore, the change in correlation between pain intensity and FC of sgACC and PAG between pain and pain-free states was significant (t=2.38; p=0.021) before, but not after BF correction. Before induction of tonic pain, pain intensity of subsequently experienced pain was positively correlated with the FC of LPBN and PAG (only before BF correction), but FC measured during the tonic pain experience was not significant (Fig. 2C). The change between states did not significantly change the relationship between the FC of LPBN and PAG and pain intensity (t=1.24; p=0.22). Before induction of tonic pain, pain intensity of subsequently experienced pain was positively correlated with the FC of RPBN and PAG, but FC measured during the tonic pain experience was not significant (Fig. 2D). The change between states significantly changed the relationship between the FC of RPBN and PAG and pain intensity (t=-3.17; p=0.0026), surviving BF correction (15 possible changes in FC).

**Table 1.**

| Functional Connectivity Dyad | State | R-score | t-stat | p-value |
| --- | --- | --- | --- | --- |
| 1. L-PBN to left amygdala | Pain-free | -0.0087 | 0.06 | 0.95 |
| 2. L-PBN to left amygdala | Pain | 0.020 | 0.14 | 0.89 |
| 3. R-PBN to right amygdala | Pain-free | 0.236 | 1.68 | 0.099 |
| 4. R-PBN to right amygdala | Pain | 0.0088 | 0.06 | 0.95 |
| 5. pACC to left amygdala | Pain-free | 0.152 | 1.07 | 0.29 |
| 6. pACC to left amygdala | Pain | -0.0836 | -0.581 | 0.56 |
| 7. pACC to right amygdala | Pain-free | 0.0302 | 0.209 | 0.84 |
| 8. pACC to right amygdala | Pain | -0.0976 | -0.679 | 0.50 |
| 9. sACC to left amygdala | Pain-free | -0.0551 | -0.382 | 0.70 |
| 10. sACC to left amygdala | Pain | -0.0146 | -0.101 | 0.92 |
| 11. sACC to right amygdala | Pain-free | -0.133 | -0.93 | 0.36 |
| 12. sACC to right amygdala | Pain | -0.0316 | -0.22 | 0.83 |
| 13. spACC to left amygdala | Pain-free | 0.0840 | 0.58 | 0.56 |
| 14. spACC to left amygdala | Pain | -0.190 | -1.34 | 0.19 |
| 15. spACC to right amygdala | Pain-free | 0.116 | 0.81 | 0.42 |
| 16. spACC to right amygdala | Pain | -0.174 | -1.22 | 0.23 |
| 17. L-PBN to PAG | Pain-free | 0.341 | 2.51 | 0.016 |
| 18. L-PBN to PAG | Pain | 0.180 | 1.27 | 0.21 |
| **19. R-PBN to PAG** | **Pain-free** | **0.444** | **3.43** | **0.001** |
| 20. R-PBN to PAG | Pain | 0.0375 | 0.26 | 0.80 |
| 21. sACC to L-PBN | Pain-free | -0.271 | -1.95 | 0.057 |
| 22. sACC to L-PBN | Pain | 0.114 | 0.79 | 0.43 |
| 23. sACC to R-PBN | Pain-free | -0.0828 | -0.58 | 0.57 |
| 24. sACC to R-PBN | Pain | -0.0133 | -0.09 | 0.93 |
| 25. pACC to PAG | Pain-free | 0.174 | 1.22 | 0.23 |
| 26. pACC to PAG | Pain | 0.0641 | 0.45 | 0.66 |
| 27. sACC to PAG | Pain-free | 0.0618 | 0.43 | 0.67 |
| **28. sACC to PAG** | **Pain** | **0.376** | **2.81** | **0.007** |
| 29. spACC to PAG | Pain-free | 0.229 | 1.63 | 0.11 |
| 30. spACC to PAG | Pain | 0.0663 | 0.46 | 0.65 |

*italics - significant before BF correction. BOLD - significant after BF correction.*

### Resting State Functional Connectivity: Seed-driven networks during pain-free and prolonged tonic pain states

Seed-driven FC from the LPBN (R>0.12) during the resting state scan with a warm thermode on the participant’s leg (hereafter, ‘pain-free state’) demonstrated significant BOLD signal correlation subcortically with the left ventral striatum, left medial dorsal region of the thalamus and right caudate nucleus (Fig. 3A and Supplemental Table 1). LPBN showed cortical FC with the left anterior midcingulate cortex (aMCC), right lingual gyrus, right medial precuneus and left parietal-occipital sulcus. After induction of heat allodynia during the tonic pain state, LPBN whole brain FC was greater on average and showed additional regions of FC (R>0.12). Additional regions of FC with LPBN included left postcentral gyrus, left ventral anterior insula, right dorsal posterior insula, bilateral inferior frontal gyri and bilateral lateral PAG (Fig. 3B and Supplemental Table 2).

**Figure 3.**
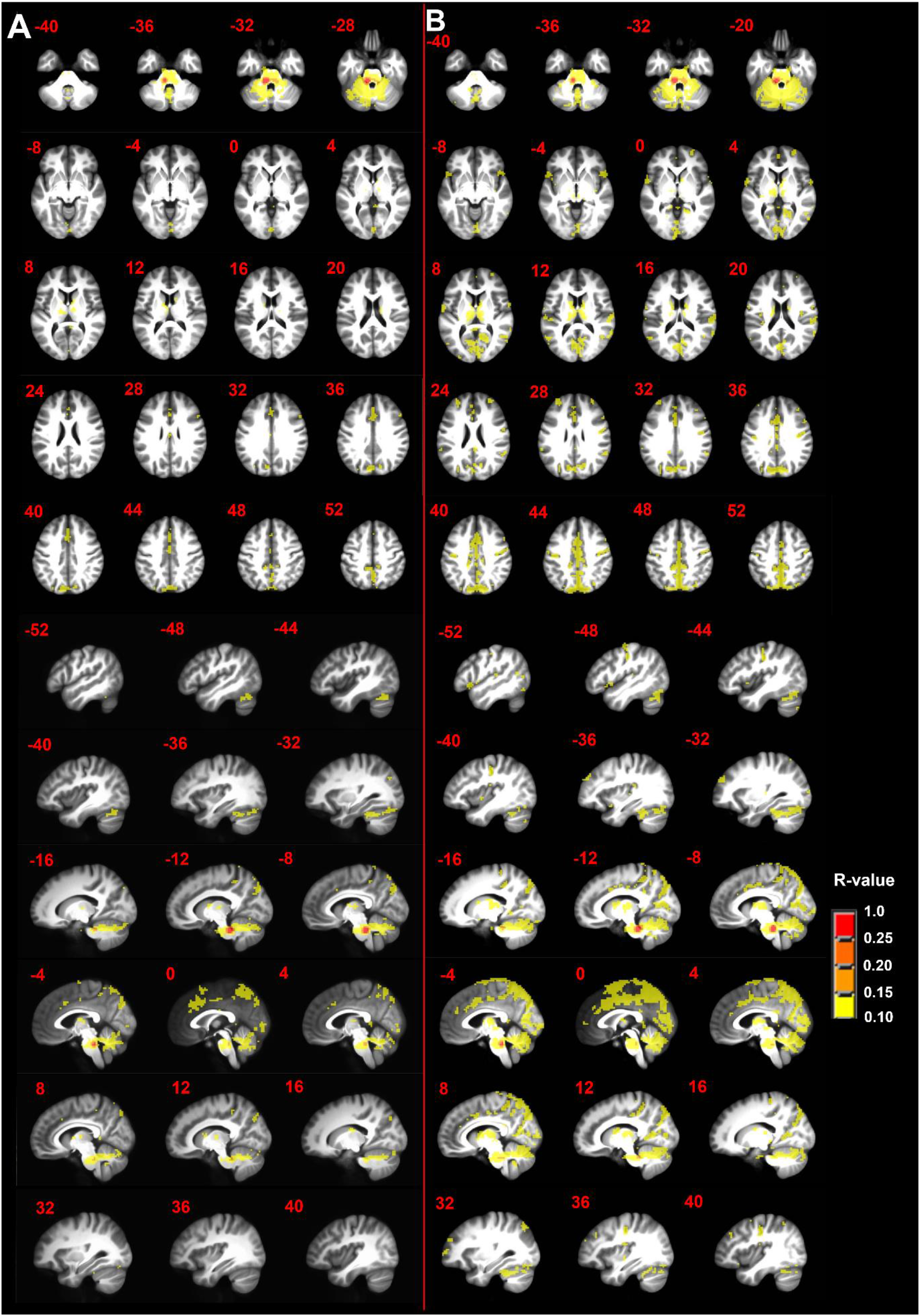
**A)** Seed-driven functional connectivity from left parabrachial complex during the pain-free state and **B)** during the tonic pain state. n=50; minimum cluster size 108 mm^3^; p-value threshold 0.0001; R-value threshold 0.10. Axial and sagittal labels are in MNI coordinates.

Seed-driven FC from the RPBN (R>0.12) during the pain-free state included subcortical regions including the contralateral LPBN, midline rostral pons, right caudate nucleus, right substantia nigra, bilateral ventral striatum, bilateral cerebellar tonsil, and left thalamus including the lateral geniculate nucleus and pulvinar (Fig. 4A and Supplemental Table 3). Pain-free state FC from the RPBN to cortical regions including left fusiform gyrus, left precentral gyrus, left middle frontal gyrus, left middle temporal gyrus, right cuneus, right inferior frontal gyrus pars orbitalis, right middle frontal gyrus, right lingual gyrus, right superior frontal gyrus, right dorsal posterior insula, bilateral transverse temporal gyri, and bilateral postcentral gyrus. During the tonic pain state FC from the RPBN (R>0.12) was greater and more extensive on average and included several new regions (Fig. 4B). New regions of FC included left inferior occipital gyrus, left inferior frontal gyrus pars orbitalis, left cuneus and calcarine gyrus, left parietal operculum, left superior frontal gyrus, right middle frontal gyrus, left inferior semilunar lobule of the cerebellum, right aMCC, right medial and lateral postcentral gyrus, right inferior frontal gyrus, right paracentral lobule, right cerebellar culmen, bilateral medial dorsal thalamus (Supplemental Table 4).

**Figure 4.**
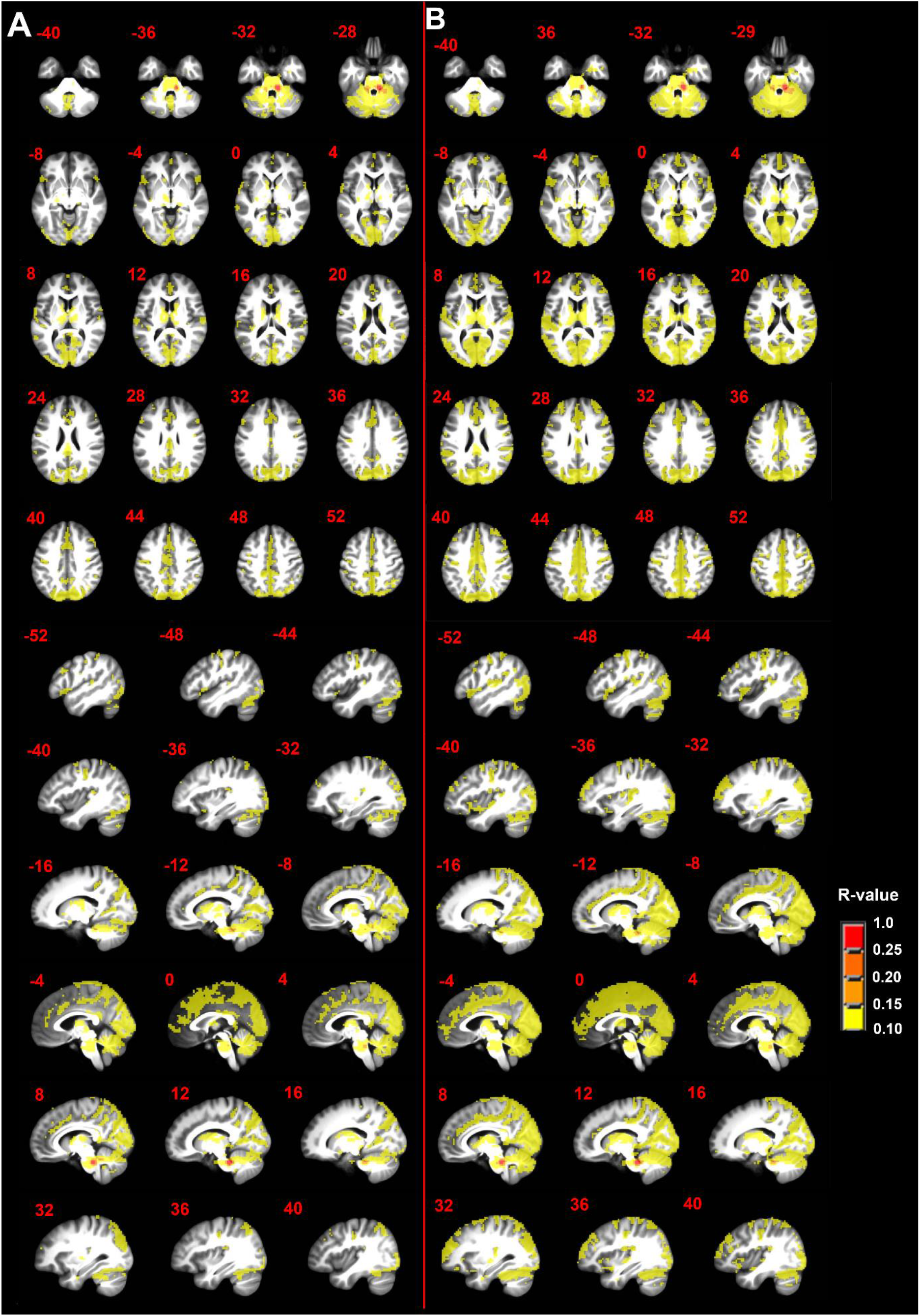
**A)** Seed-driven functional connectivity from right parabrachial complex during the pain-free state and **B)** during the tonic pain state. n=50; minimum cluster size 108 mm^3^; p-value threshold 0.0001; R-value threshold 0.10. Axial and sagittal labels are in MNI coordinates.

PAG seed-driven FC (R>0.12) during the pain-free state was consistent with previously published FCNs of the PAG including the sgACC, medial thalamus, bilateral lingual gyri, bilateral cerebellar declive and bilateral caudal pons (Supplemental fig. 1A, Supplemental Table 5) (Kong et al., 2010; Yu et al., 2014). During the tonic pain state PAG seed-driven FC included additional brain regions including aMCC, left paracentral lobule, left postcentral gyrus and rostral ventromedial medulla (Supplemental fig. 1B, Supplemental Table 6).

Seed-driven FC during the pain-free state from the LAMYG (R>0.20) was consistent with previous reports including significant FC in the bilateral ventromedial prefrontal cortex, frontal operculum, paracentral lobule, fusiform gyri, postcentral gyri, precentral gyri, hippocampus, dorsal posterior and ventral anterior insula, contralateral (right) amygdala and ipsilateral (left) temporo-occipital junction (Supplemental fig. 2A, Supplemental Table 7) (Gorka et al., 2018; Kerestes et al., 2017; Roy et al., 2009; Simons et al., 2014). During the tonic pain state, FC was stronger and more extensive from the LAMYG seed to the midcingulate cortex, bilateral anterior and posterior insula, bilateral inferior frontal gyri pars orbitalis, bilateral posterior cingulate cortex, left superior parietal lobule and right superior temporal gyrus (Supplemental fig. 2B, Supplemental Table 8).

Seed-driven FC during the pain-free state from the RAMYG (R>0.20) was consistent with previous reports and that driven from the LAMYG, excepting additional significant FC in the bilateral superior temporal gyri, temporoparietal junction, middle temporal gyri, contralateral (left) amygdala and selective FC limited to the ipsilateral (right) dorsal posterior insula (Supplemental fig. 3A, Supplemental Table 9). During the tonic pain state, FC was stronger and more extensive from the RAMYG seed to the bilateral posterior insula, left paracentral lobule, left inferior parietal lobule and right fusiform gyrus (Supplemental fig. 3B, Supplemental Table 10).

### Resting State Functional Connectivity: Brain-wide contrast between pain and pain-free states

To determine the statistically significant difference between the seed-derived FC of the resting state maps collected during the pain-free task and prolonged tonic pain, we calculated brain-wide contrast maps (Figure 5). After cluster extent correction at peak p-value of 0.001, there were no significant contrast results from the PAG or RAMYG seeds. During the tonic pain state, the FC from the LAMYG was significantly greater in left middle frontal gyrus (BA8), left superior frontal gyrus, right precentral gyrus, right middle frontal gyrus and bilateral precuneus (Fig. 5A, Table 2). During the tonic pain state, the FC from the LPBN was significantly greater in the right precentral gyrus (volume=270 mm^3^, max t-stat=4.48, MNI coordinates: x=44, y=-14, z=47) (Fig. 5B). During the tonic pain state, the FC from the RPBN was significantly greater in bilateral medial postcentral gyrus (Fig. 5C, Table 3).

**Figure 5.**
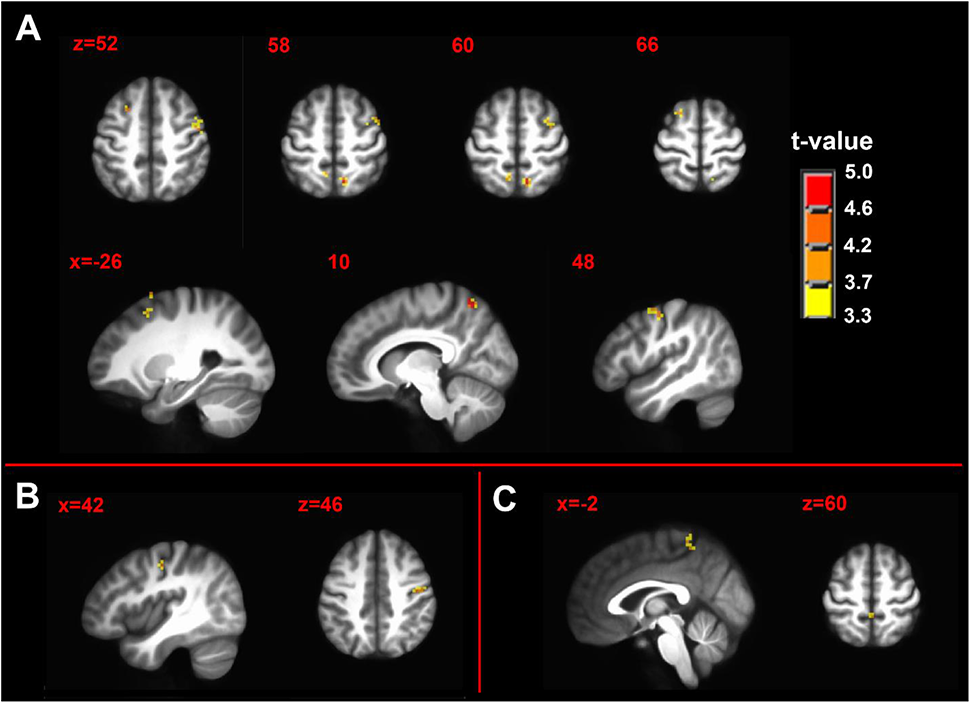
Contrast maps of tonic pain state > pain-free state of seed-driven functional connectivity from **A)** left amygdala complex seed **B)** left parabrachial complex seed and **C)** right parabrachial complex seed. n=50; minimum cluster size 270 mm^3^; p-value threshold 0.001. Axial and sagittal labels are in MNI coordinates.

**Table 2.** Seed: Left Amygdala Contrast: Pain > Control (p<0.001)

| Cluster Number and Brain Region | Brodmann Area | Volume | Maximum Intensity | MNI Coordinates |
| --- | --- | --- | --- | --- |
| 1. Right Precentral Gyrus | 4 | 810 | 4.70 | (52, -11, 50) |
| 2. Right Precuneus | 7 | 567 | 5.00 | (11, -62, 62) |
| 3. Left Middle Frontal Gyrus | 8 | 324 | 4.14 | (-23, 14, 53) |
| 4. Left Precuneus | 7 | 270 | 4.06 | (-8, -56, 62) |
| 5. Right Middle Frontal Gyrus | 6 | 270 | 4.01 | (35, -5, 62) |
| 6. Left Superior Frontal Gyrus | 6 | 270 | 4.07 | (-20, 14, 71) |

**Table 3.** Seed: Right Parabrachial Nucleus Contrast: Pain > Control (p<0.001)

| Cluster Number and Brain Region | Brodmann Area | Volume | Maximum Intensity | MNI Coordinates |
| --- | --- | --- | --- | --- |
| 1. Right Postcentral Gyrus | 5 | 513 | 4.84 | (11, -44, 71) |
| 2. Left Postcentral Gyrus | 5 | 459 | 4.31 | (-5, -44, 68) |

### Resting State Functional Connectivity: Pain intensity covariance with seed-based FC

In the pain intensity covariation analysis, after cluster extent correction at peak p-value of 0.005, there were no significant correlations of pain intensity with FC from left or right parabrachial nuclei during pain-free or tonic pain states.

Pain intensity was significantly positively correlated with FC between the LAMYG seed during the pain-free state and right superior parietal lobule, right calcarine gyrus and right caudate nucleus (Fig. 6A, Table 4). During the tonic pain state, pain intensity was significantly negatively correlated with FC between the LAMYG seed and right inferior temporal gyrus and right superior frontal gyrus (Fig. 6B, Table 5). Partial correlation analysis demonstrated no interdependence among seed regions and significant clusters for the LAMYG during the pain-free or tonic pain states.

**Figure 6.**
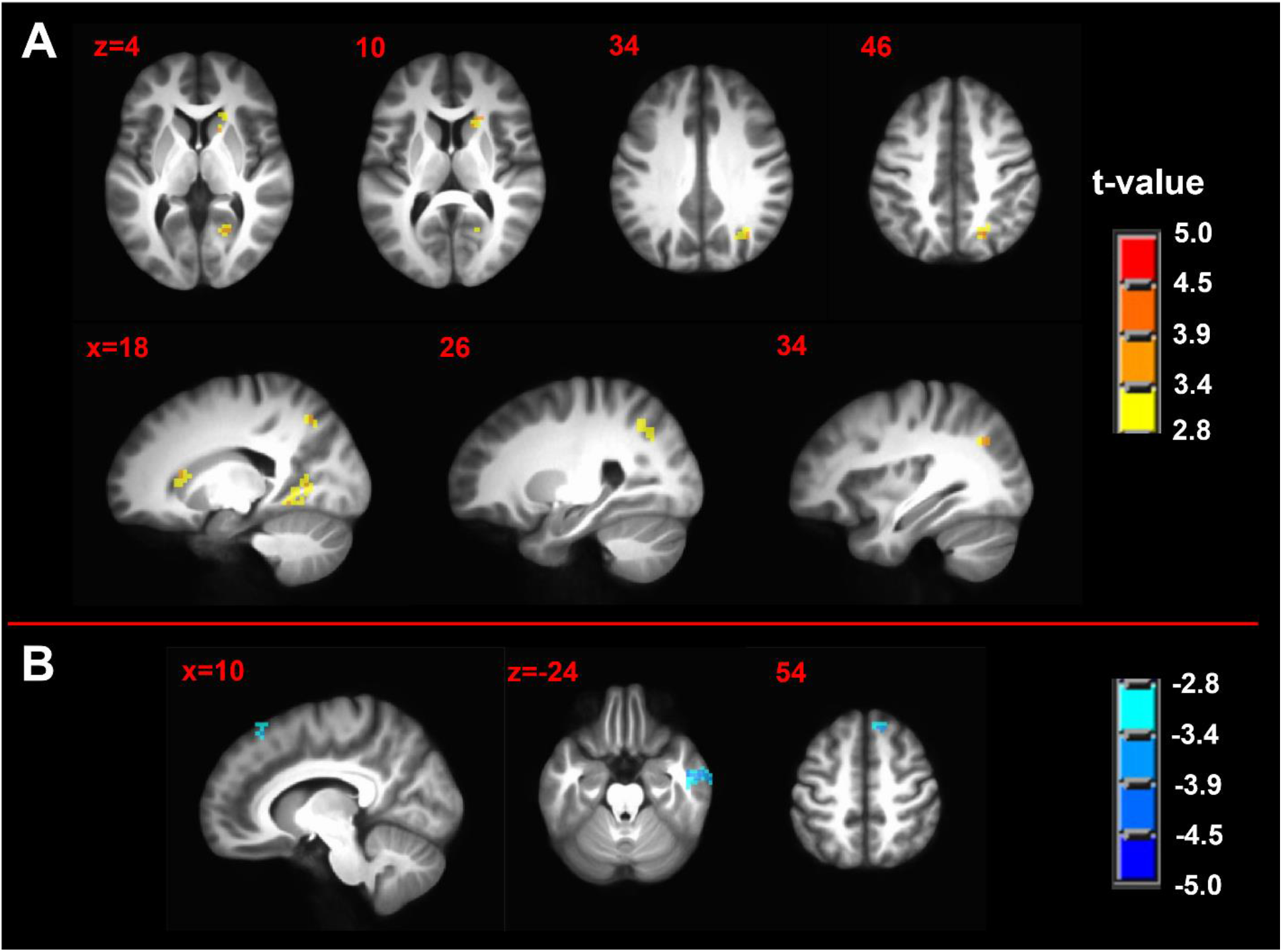
Pain intensity covariation with seed-driven functional connectivity maps of **A)** left amygdala complex seed during pain-free state immediately before **B)** left amygdala complex seed during tonic pain state. n=50; minimum cluster size 540 mm^3^; p-value threshold 0.005. Axial and sagittal labels are in MNI coordinates.

**Table 4.** Seed: Left Amygdala State: Control Test: Correlation with Pain Intensity ( $p < 0.005$ )

| Cluster Number and Brain Region | Brodmann Area | Volume | Maximum Intensity | MNI Coordinates |
| --- | --- | --- | --- | --- |
| 1. Right Superior Parietal Lobule | 7 | 1161 | 3.96 | (23, -65, 50) |
| 2. Right Calcarine Gyrus | 17 | 918 | 3.62 | (23, -62, 5) |
| 3. Right Caudate Nucleus |  | 729 | 3.80 | (20, 23, 14) |

**Table 5.** Seed: Left Amygdala State: Pain Test: Correlation with Pain Intensity ( $p < 0.005$ )

| Cluster Number and Brain Region | Brodmann Area | Volume | Maximum Intensity | MNI Coordinates |
| --- | --- | --- | --- | --- |
| 1. Right Inferior Temporal Gyrus | 20 | 1242 | -4.09 | (59, -8, -23) |
| 2. Right Superior Frontal Gyrus | 8 | 567 | -4.16 | (17, 32, 53) |

Pain intensity was significantly positively correlated with FC between the RAMYG seed during the pain-free state and the right claustrum, right middle temporal gyrus, right temporo-occipitoparietal junction (TOPJ) and bilateral calcarine gyri (Fig. 7A, Table 6). Partial correlation analysis revealed interdependencies between the right middle temporal gyrus and right temporo-occipitoparietal junction (Fig. 7A). Further interdependencies were revealed among left and right calcarine gyri and right claustrum (Fig 7A). Positive correlations between pain intensity during the tonic pain state and FC during the pain-free state between the RAMYG seed and significant clusters in a brain-wide analysis were R>0.51 (Fig. 7B). In contrast, there were no regions of significant correlation between pain intensity and FC between RAMYG and the brain during the tonic pain state scan.

**Figure 7.**
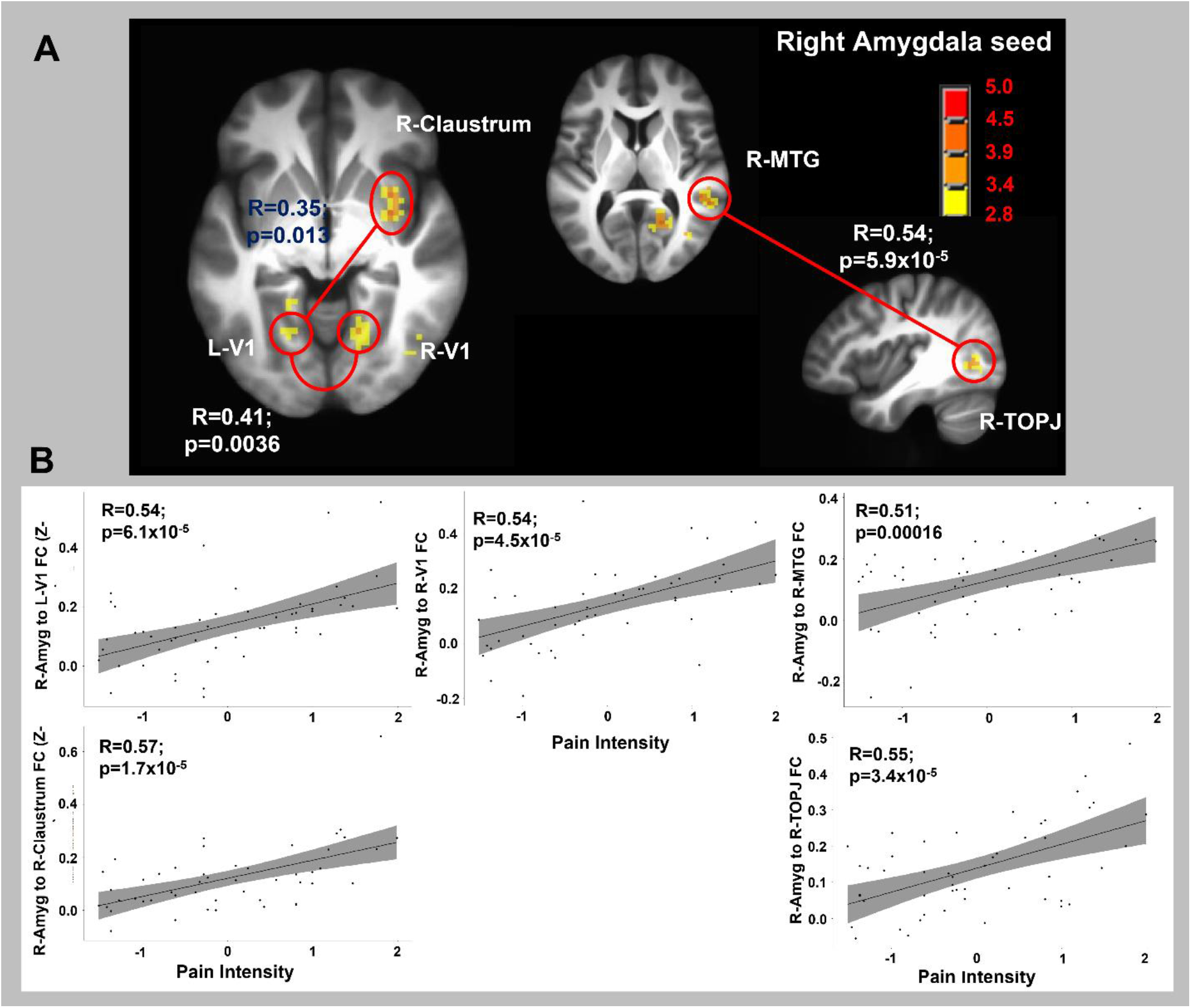
**A)** Pain intensity covariation with seed-driven functional connectivity maps of right amygdala complex seed during pain-free state. Display of significant partial correlation interactions from all seed partial correlation analysis. n=50; minimum cluster size 540 mm^3^; p-value threshold 0.005. **B)** Display of the positive correlation between right amygdala complex seed and significant cluster areas and pain intensity. Shaded area corresponds to the area of 95% confidence interval.

**Table 6.** Seed: Right Amygdala State: Control Test: Correlation with Pain Intensity ( $p < 0.005$ )

| Cluster Number and Brain Region | Brodmann Area | Volume | Maximum Intensity | MNI Coordinates |
| --- | --- | --- | --- | --- |
| 1. Right Calcarine Gyrus | 17 | 2943 | 4.08 | (20, -59, 5) |
| 2. Right Claustrum |  | 837 | 4.30 | (32, 8, -5) |
| 3. Right Middle Temporal Gyrus | 37 | 756 | 3.78 | (44, -71, -2) |
| 4. Right Temporo-occipitoparietal Junction |  | 756 | 4.15 | (50, -38, 8) |
| 5. Left Calcarine Gyrus | 17 | 675 | 4.01 | (-20, -47, -8) |

Pain intensity was significantly positively correlated with FC between the PAG seed during the pain-free state and left cerebellum, left middle frontal gyrus, right posterior cingulate cortex and paracentral lobule, right inferior parietal lobule, right medial precuneus and RPBN (Fig. 8A, Table 7). Partial correlation analysis revealed a complex web of interdependencies among the identified clusters (Fig. 8A). Positive correlations between pain intensity during the tonic pain state FC during the pain-free state between the PAG seed and significant clusters in a brain-wide analysis were R>0.49 (Fig. 8B). In contrast, there were no regions of significant correlation between pain intensity and FC between PAG and the whole brain during the tonic pain state.

**Figure 8.**
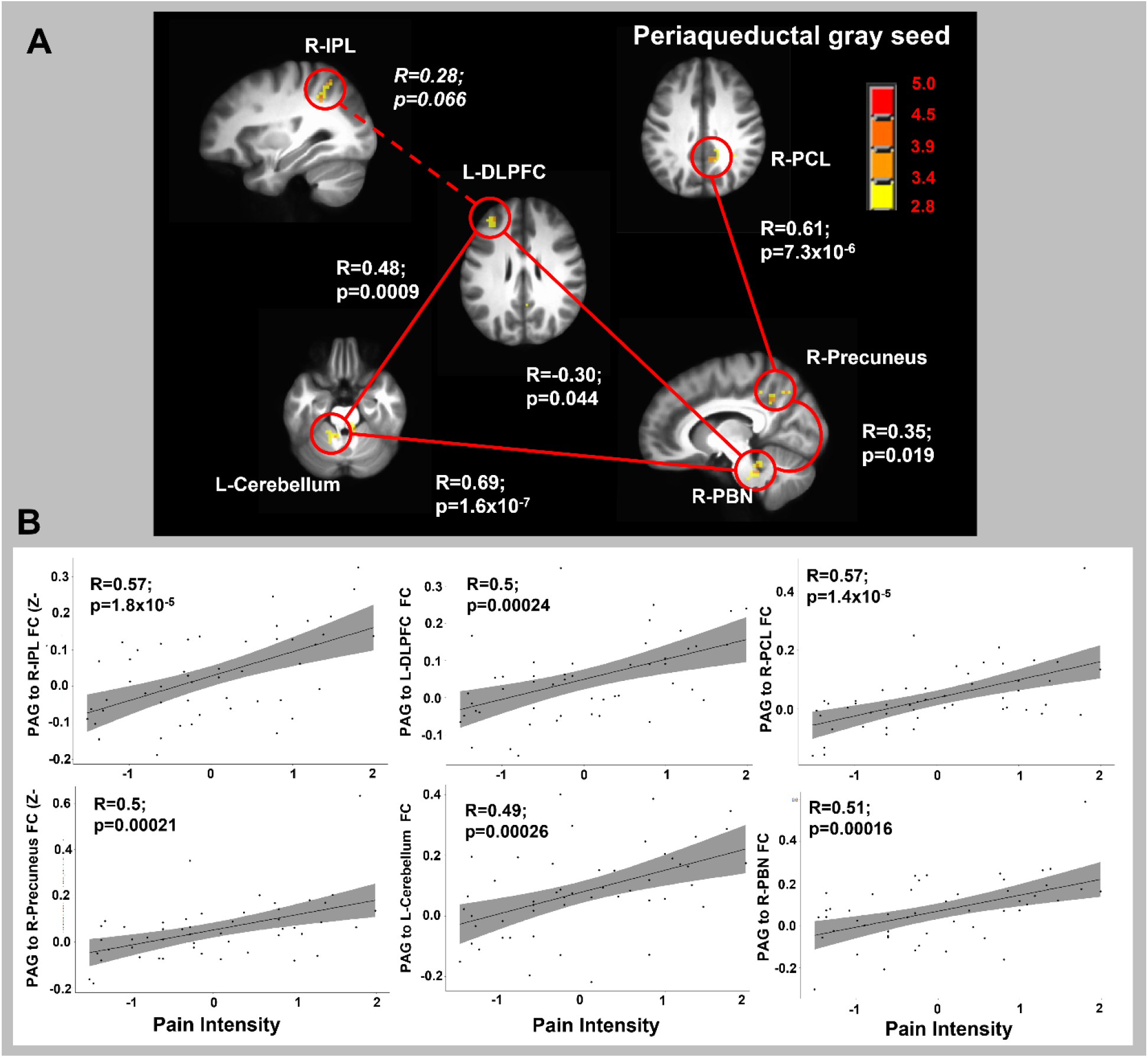
**A)** Pain intensity covariation with seed-driven functional connectivity maps of periaqueductal gray area seed during pain-free state. Display of significant partial correlation interactions from all seed partial correlation analysis. Dotted line represents a connection that did not quite reach traditional statistical significance. n=50; minimum cluster size 540 mm^3^; p-value threshold 0.005. **B)** Display of the positive correlation between right amygdala complex seed and significant cluster areas and pain intensity. Shaded area corresponds to the area of 95% confidence interval.

**Table 7.**
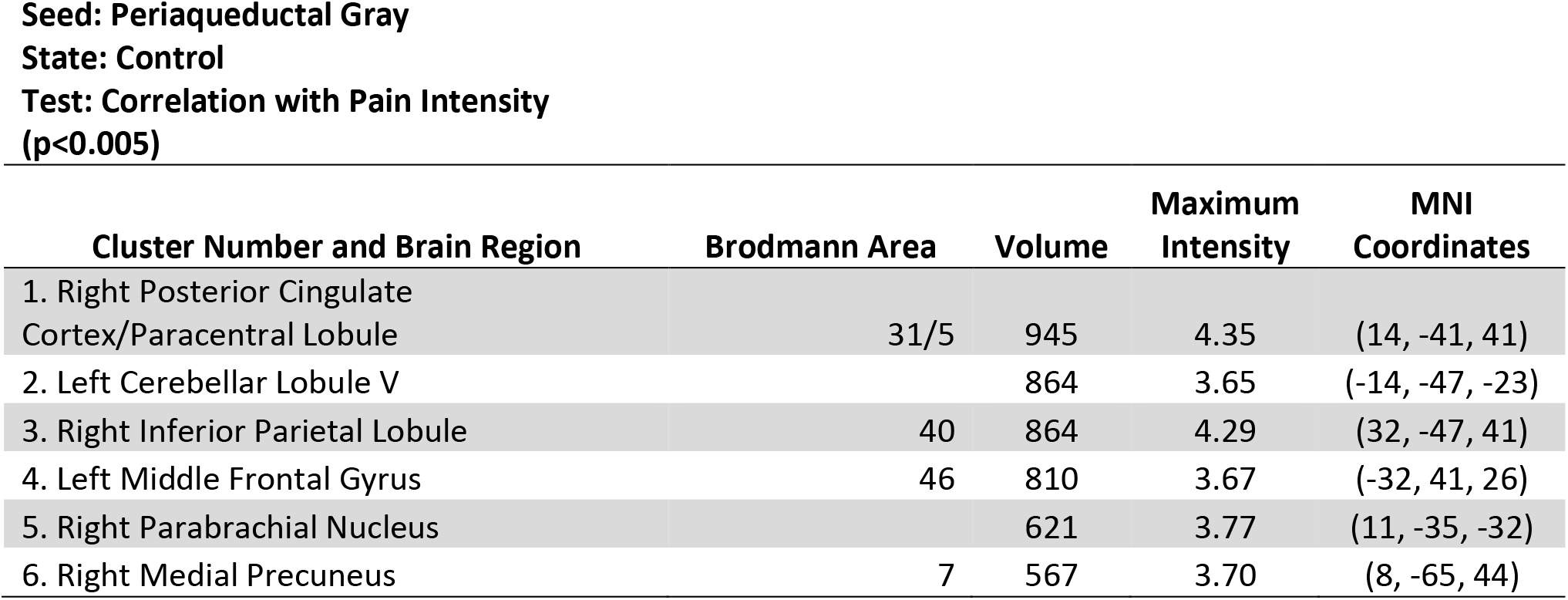

## Discussion

The purpose of this study was to produce tonic pain via a reliable, interindividually variable model to investigate resting state FC of the DPMN in subcortical brain areas. We operationalized this by hypothesizing specific modulation of FC between several FC dyads including spACC-PAG, pACC-PAG, sgACC-PAG, sgACC-LPBN, sgACC-RPBN, LPBN-PAG, RPBN-PAG, LAMYG-LPBN, and RAMYG-RPBN. Given our limited dyadic FC changes during tonic pain in pain-free participants, we completed exploratory analyses including tests for brain-wide changes in FC during pain-free and tonic pain states. Finally, we conducted exploratory analysis covarying pain intensity with brain-wide FC.

### Limited evidence of disruption of node-to-node FC in DPMN

Suprathreshold pain in the C-HP model was robust and highly variable between individuals, inducing robust heat allodynia for at least 75 minutes (Fig. 1B). Within our hypothesized group of 9 dyads there was only evidence of pain disrupting FC between sgACC-PAG (Fig. 2A). In contrast, the effect of sex demonstrated greater pACC-PAG FC in males than females, as previously found in PAG seed-driven FC analyses (Karshikoff et al., 2016). Prior reports of sex differences in FC between ACC and PAG was the driver behind including sex as a factor in our analysis (Tomasi and Volkow, 2012; Wang et al., 2014). For example, increases in brain-wide PAG FC in response to phasic pain are greater in males than female, including with LAMYG. Given the relatively large sample size in this study, failure to find effects of pain in these subcortical connections suggests a relatively weak contribution to changing perception of warmth to pain.

### FC within the DPMN can predict pain intensity during tonic pain

While the sgACC-PAG FC during pain-free rest was unrelated to future pain intensity, it became positively related to pain intensity during tonic pain. Both left and right PBN FC to PAG during pain-free rest were positively related to subsequent pain, but during tonic pain this relationship was abolished (Fig. 2C and 2D). PBN FC with PAG at rest predicting pain intensity is strongly supportive of the involvement of this connection in pain regulation in humans, consistent with recent findings in rodent models (Huang et al., 2019; Raver et al., 2020; Roeder et al., 2016).

Our results demonstrate FC disruption between sgACC-PAG by tonic pain, and the first evidence of PBN-PAG FC predicting to greater tonic pain experience in humans. It should be noted that our correlational results may support stronger inferences relative to state change results, since our participants experienced pain intensity ranging from just above threshold to severe (75/100 NRS). This variability in pain experience was present with lower exposure temperatures for those participants with greater reported pain intensity and higher exposure temperatures for participants with lower reported pain intensity. This suggests a sensitivity measure of the model would demonstrate much greater effect variability compared to pain intensity.

FC and BOLD fMRI modulations associated with pain modulation often include effects in the ACC and PAG. The regions of ACC involved in pain modulation effects associated with PAG include 1) pACC (Coulombe et al., 2016; Eippert et al., 2009; Harper et al., 2018; Kong et al., 2010; Labus et al., 2013; Lee et al., 2021; Leknes et al., 2013; Li et al., 2016; Linnman et al., 2012; Meeker et al., 2019; Pecina et al., 2015; Peyron et al., 2007; Schmidt-Wilcke et al., 2014; Valet et al., 2004; Wager et al., 2007; Wagner et al., 2001; Wey et al., 2014; Yu et al., 2017; Yu et al., 2014), 2) sgACC (Ayoub et al., 2021; Bingel et al., 2006; Labus et al., 2013; Lee et al., 2021; Leknes et al., 2013; Li et al., 2020; Meeker et al., 2019; Pecina et al., 2015; Schmidt-Wilcke et al., 2014; Sprenger et al., 2011; Wang et al., 2014) or 3) spACC (Atlas et al., 2012; Coulombe et al., 2016; Kong et al., 2010; Lee et al., 2021; Linnman et al., 2012; Petrovic et al., 2002; Peyron et al., 2007; Rezai et al., 1999; Schmidt-Wilcke et al., 2014; Schulz et al., 2020; Wager et al., 2007; Wagner et al., 2001). Complementarily, non-human primate tractography studies demonstrate afferents projecting to the PAG from pACC and spACC, aMCC, and sgACC without reciprocal connections from PAG to cortex (An et al., 1998; Mantyh, 1982b; Porrino and Goldman-Rakic, 1982; Vogt et al., 1987).

### Anatomical evidence of PBN connectivity in animal models and humans

An important spinocortical pathway for nociceptive information from the periphery, most clearly delineated in rodents, that communicates with the DPMN is the spinoparabrachial pathway (Bernard and Besson, 1990; Gauriau and Bernard, 2002; Polgar et al., 2010; Raver et al., 2020; Roeder et al., 2016). This pathway bypasses the thalamus to project to the cortex directly or via the amygdala and is sensitized by prolonged tonic thermal stimuli (Kissiwaa and Bagley, 2018). In primates, the lateral PBN projects to central nucleus of the amygdala, bed nucleus of the stria terminalis, ventroposteromedial, central lateral, parafascicular/centromedian and reuniens nuclei of the thalamus; dorsomedial, lateral, ventromedial, supramamillary and infundibular nuclei of the hypothalamus; the midbrain PAG, substantia nigra and ventral tegmental area; medullary nucleus ambiguous and reticular formation; and receives a projection from the sgACC (Freedman et al., 2000; Pritchard et al., 2000). The primate medial PBN projects to frontal polar cortex, the lateral principal sulcus, medial BA9; and pACC and spACC of the cortex (Porrino and Goldman-Rakic, 1982). Projections from the medial PBN reach the central nucleus of the amygdala; bed nucleus of the stria terminalis; the lateral, dorsal and dorsomedial nuclei of the hypothalamus; ventroposteromedial, central lateral, parafascicular/centromedian and reuniens nuclei of the thalamus; and midbrain PAG and annular nucleus (Pritchard et al., 2000). The lateral and medial parabrachial nuclei are separated by the brachium conjunctivum (Porrino and Goldman-Rakic, 1982). We did not distinguish between lateral and medial PBN in our seeds since we are constrained by low functional resolution and partial volume effects. Relatively few human neuroimaging of studies of the neural correlates of painful stimuli have previously reported PBN activation (Dunckley et al., 2005; Fairhurst et al., 2007; Sprenger et al., 2011; Stroman et al., 2018).

In our FC study of the PBN, we found significant BOLD FC with the caudate, ventral striatum, and medial thalamus subcortically (Fig. 3 and 4). Cortically, we found significant BOLD FC with lingual and fusiform gyri, anterior MCC, precentral gyri, transverse temporal gyri, postcentral gyri and anterior and posterior insula. Many of these brain areas likely represent polysynaptic influences of pontine parabrachial complex activity and may aid the PBN complex’s role in interoception, pain processing, regulation of food intake and thermoregulation (Benarroch, 2016). Almost the entire pons was included in the cluster of significant FC along with the seed region, obscuring brainstem regions involved in the PBN complex’s role in arousal and respiratory control. Importantly, PBN complex FC included areas involved in processing sensory (e.g., primary sensory, posterior insula), affective (e.g., anterior MCC, anterior insula and medial thalamus), and motivational (e.g., BA46, BA9) aspects of pain processing (Kulkarni et al., 2005; Melzack and Casey, 1968; Rainville et al., 1999a). This multimodal processing capability is consistent with animal studies combining ascending and descending pain processing regions extending rodent model findings to a possible role in pain processing and modulation in humans (Chen and Heinricher, 2019; Kissiwaa and Bagley, 2018; Raver et al., 2020; Roeder et al., 2016; Uddin et al., 2018).

### Modulation of subcortical FC with cortical motor and sensory processing areas

Contrasts between pain-free and pain states of LPBN FC showed enhanced FC to the lateral precentral gyrus in BA4, and from RPBN showed enhanced FC to the medial crown of the postcentral gyrus (Fig 5B and 5C). The bilateral medial postcentral gyrus area corresponds to the leg representation of somatosensory cortex, where the painful stimulus was applied. The anterior lateral motor cortex area may be involved in a greater eye-movement or blinking between pain and pain-free states (Eisenach et al., 2017; Paparella et al., 2020). The LAMYG to whole brain FC contrast demonstrated enhanced FC to the bilateral precuneus, right precentral gyrus, and left middle and superior frontal gyri (Fig. 5A). The significant areas of the BOLD FC changes in the precuneus were in the sensorimotor section with projections to frontal motor control, sensory parietal, and paracentral areas of the brain (Margulies et al., 2009; Morecraft et al., 2004). Enhanced FC from sensorimotor representations of affected body sites to AMYG and PAG replicate FC aberrations found in chronic low back pain, chronic neck pain and pressure pain in pain-free participants (Kim et al., 2015; Kim et al., 2013; Kong et al., 2013; Yu et al., 2017; Yu et al., 2014).

Given the variability of reported tonic pain intensity in the MRI scanner environment (range 1-75 out of 100 NRS), we looked for regions of brain-wide FC correlated with pain intensity either during pain-free rest or during tonic pain. During pain-free rest, BOLD FC emanating from the LAMYG to right caudate, right superior parietal lobule and right calcarine gyrus positively correlated with subsequent pain (Fig. 6A). This implies cross-hemisphere amygdala FC to higher order sensory cortices is stronger in those individuals more sensitive to pain. This may be related to individual variability in negative affect as recent evidence in pain-free participants showed stronger connectivity of the AMYG to S1/M1, S2/operculum, and posterior parietal cortex at rest in individuals with greater pain facilitation by negative emotions (Gandhi et al., 2020).

### Extensive amygdalocortical and PAG-cortical FC predicts pain intensity during tonic pain

Seed-driven FC from RAMYG to the whole brain positively predicted pain intensity experienced during tonic pain specifically in the bilateral visual cortex, right middle temporal gyrus, right temporo-occipital temporal gyrus and left claustrum (Fig. 7). Partial correlation analyses revealed two networks emanating from the RAMYG that predicted tonic pain including 1) RAMYG, right claustrum, and left and right primary visual cortex; and 2) RAMYG, right middle temporal gyrus and right temporo-occipital temporal gyrus. Brain areas in these networks are all highly connected with each other and the rest of the brain, excepting the primary visual cortex (Gorka et al., 2018; Heilbronner and Haber, 2014; Kaas, 2012; Krimmel et al., 2019; Miyashita et al., 2007). While it may be surprising to observe primary visual cortex implicated in a network correlated with pain intensity, a possible ‘vascular steal’ effect resulting in decreases in BOLD or blood oxygenation measured by positron emission tomography, in the primary visual cortex by painful stimuli, as well as multisensory processing within the primate visual cortex both have well-established precedent (Coghill et al., 1999; Moulton et al., 2005; Mouraux et al., 2011; Murray et al., 2016).

Finally, we found seed-driven FC from PAG positively predicted pain intensity experienced during tonic pain in right paracentral lobule, RPBN, right medial precuneus, right IPL, left cerebellum, left DLPFC (Fig. 8). These regions were interconnected within a PAG-anchored network revealed through partial correlation modeling. Both left DLPFC and right IPL have been implicated in modified pain states, chronic pain states and prediction of chronic pain (Niu et al., 2019; Rainville et al., 1999b; Seminowicz et al., 2018; Seminowicz and Moayedi, 2017; Seminowicz et al., 2011; Symonds et al., 2006). The right paracentral lobule cluster was near the sensory representation of the stimulated left lower leg. Modulation of FC in sensorimotor networks to brainstem and AMYG replicates results in both healthy participants experiencing tonic pain and in chronic pain patients (Hubbard et al., 2014; Kim et al., 2015; Kim et al., 2013; Kong et al., 2013; Yu et al., 2014).

### Limitations

It is necessary to point out several limitations of our study. Some of these limitations apply broadly to resting state FC studies, while others are specific. Specific to this study, we combined resting state BOLD fMRI datasets acquired with different voxel resolutions (3 mm isotropic vs. 1.8×1.8×4 mm), did not use white matter signal as a baseline covariate, and used a 4 mm FWHM smoothing kernel. These modifications, some of which were rationalized by adjusting to use of midbrain and pontine seeds, would only decrease the signal-to-noise ratio and, given the sample size, more likely favor the presence of false negatives over false positives taking into account within subject changes (Beissner et al., 2014; Dansereau et al., 2017). Using only motion parameters and motion derivatives regression may be an issue of systematic bias, except that motion outliers, in terms of framewise displacement, were removed; and average framewise displacement was not significantly different between the pain-free and pain state (Parkes et al., 2018). We decided to retain global signal, since global signal regression has biasing effects on long-range FC and introduces negative correlation (Parkes et al., 2018; Saad et al., 2012). Furthermore, non-exploratory primary hypotheses were tested using either manually or atlas-based ROIs or spheres. Since all comparisons are within subject (except for male vs. female contrasts), significant effects were found with all biases being equal. Finally, we clearly did not evaluate all regions or connections which fit our criteria for inclusion in the DPMN as we defined it. We intend to investigate additional regions (e.g., cortical and basal ganglia seeds) in subsequent reports.

## Conclusion

The described pain model induces robust, variable, thermal allodynia allowing a sustained painful state lasting more than 45 minutes ((Anderson et al., 2002; Meeker et al., 2019) and Fig. 1). We demonstrate 1) disruption of sgACC-PAG FC during tonic pain in pain-free healthy participants; 2) sgACC-PAG FC during tonic pain positively correlates with self-report pain intensity; 3) RPBN-PAG FC during pain-free rest predicts subsequent pain intensity during tonic pain and this FC is abolished by tonic pain. Finally, we revealed brain-wide PAG- and RAMYG-anchored networks which predict pain intensity during tonic pain. Resting state FC of subcortical DPMN-associated structures such the PAG, AMYG and PBN with their cortical targets, as described in this and other recent work, may allow development of novel invasive and non-invasive neuromodulation methods and tracking of objective brain-derived biomarkers in future novel analgesic interventions. It is especially relevant to observe these findings in a large sample size of participants in a pathologically relevant human model of nociceptive central sensitization (Lotsch et al., 2014; Meeker et al., 2019; Meeker et al., 2021; Quesada et al., 2021; Simone et al., 1991).

## Supplemental Information

### Supplemental Figures

**Supplemental Figure 1.**
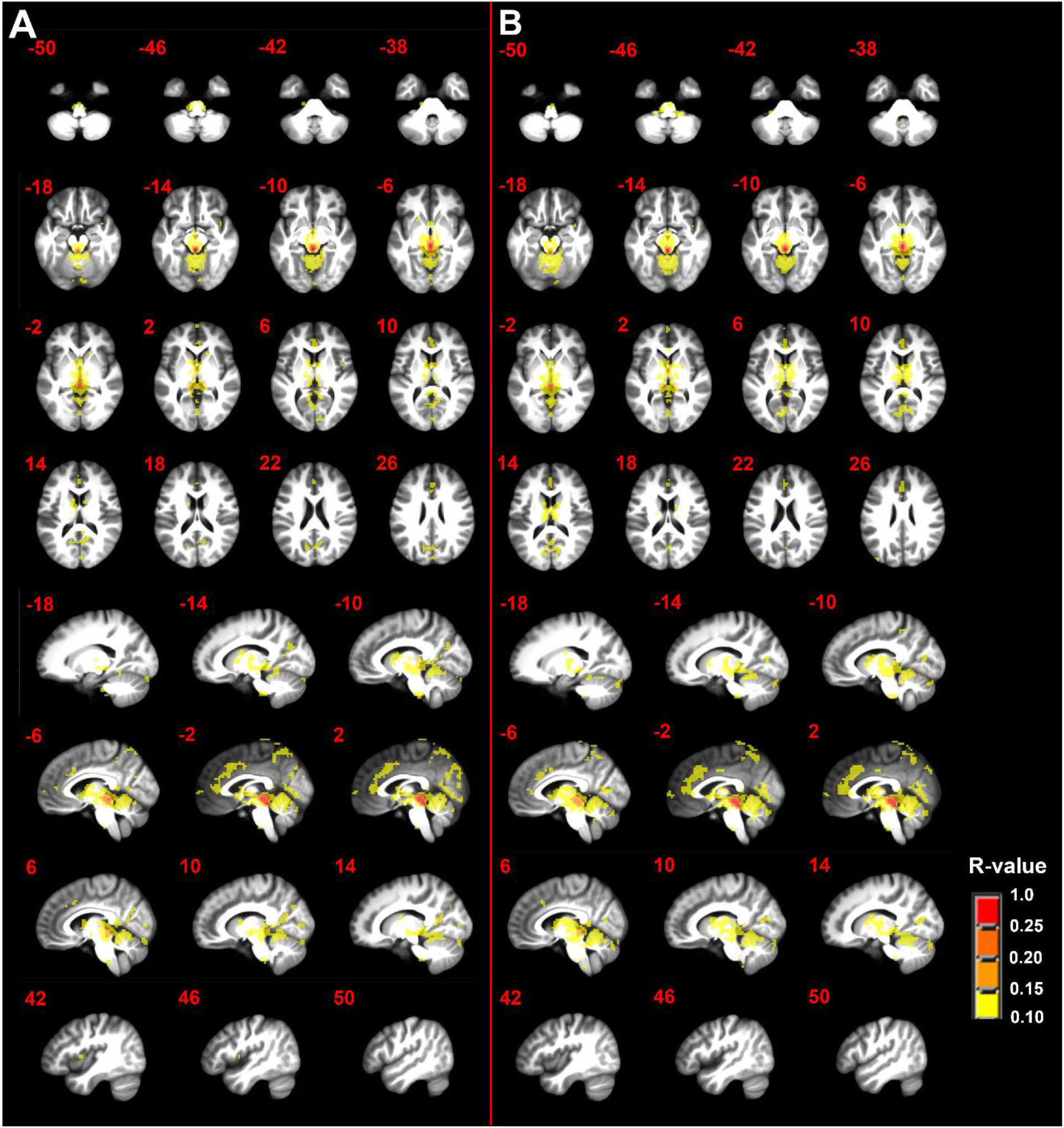
**A)** Seed-driven functional connectivity from the periaqueductal gray area during the pain-free state and **B)** during the tonic pain state. n=50; minimum cluster size 108 mm^3^; p-value threshold 0.0001; R-value threshold 0.10. Axial and sagittal labels are in MNI coordinates.

**Supplemental Figure 2.**
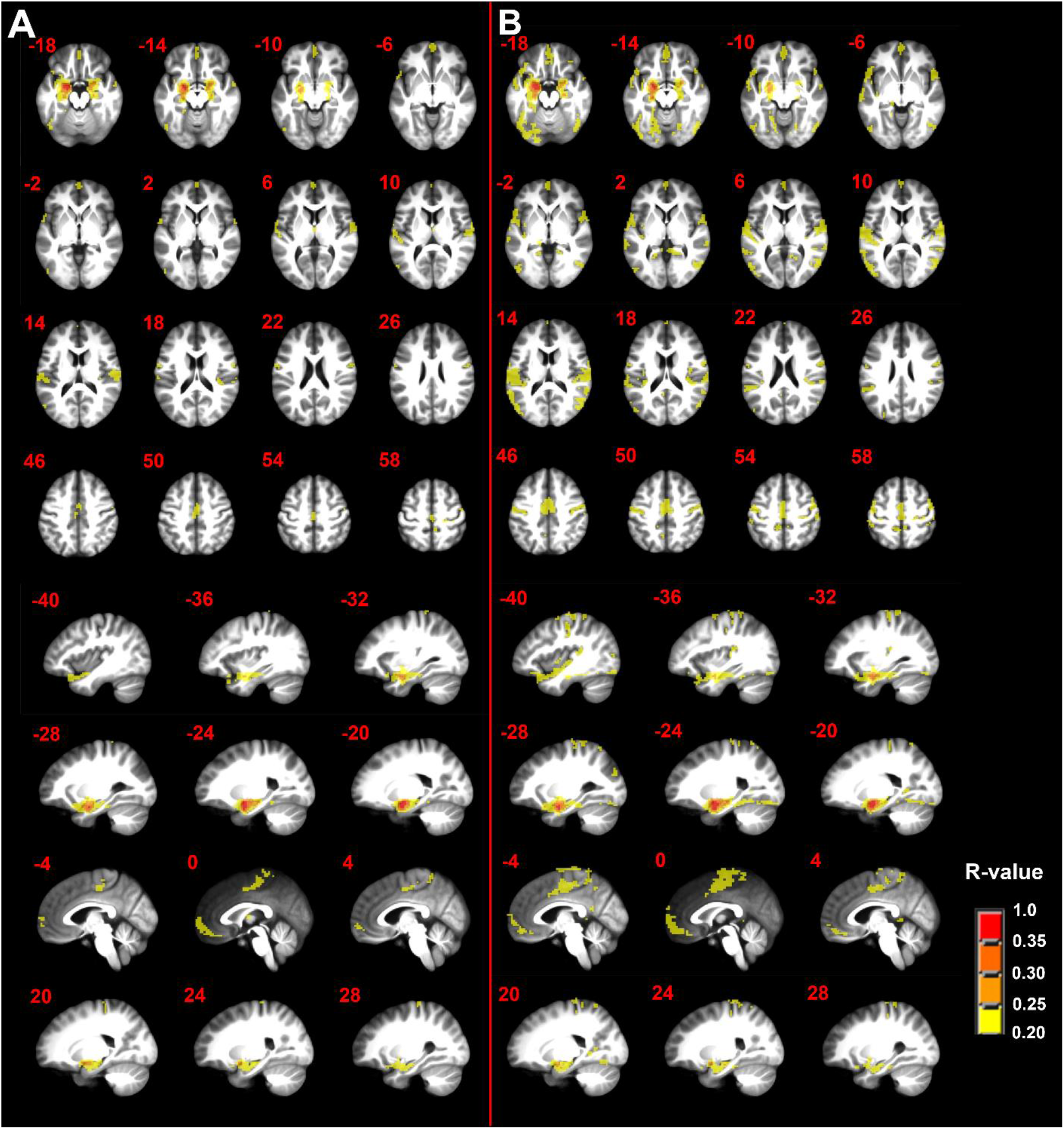
**A)** Seed-driven functional connectivity from left amygdala complex during the pain-free state and **B)** during the tonic pain state. n=50; minimum cluster size 108 mm^3^; p-value threshold 0.0001; R-value threshold 0.20. Axial and sagittal labels are in MNI coordinates.

**Supplemental Figure 3.**
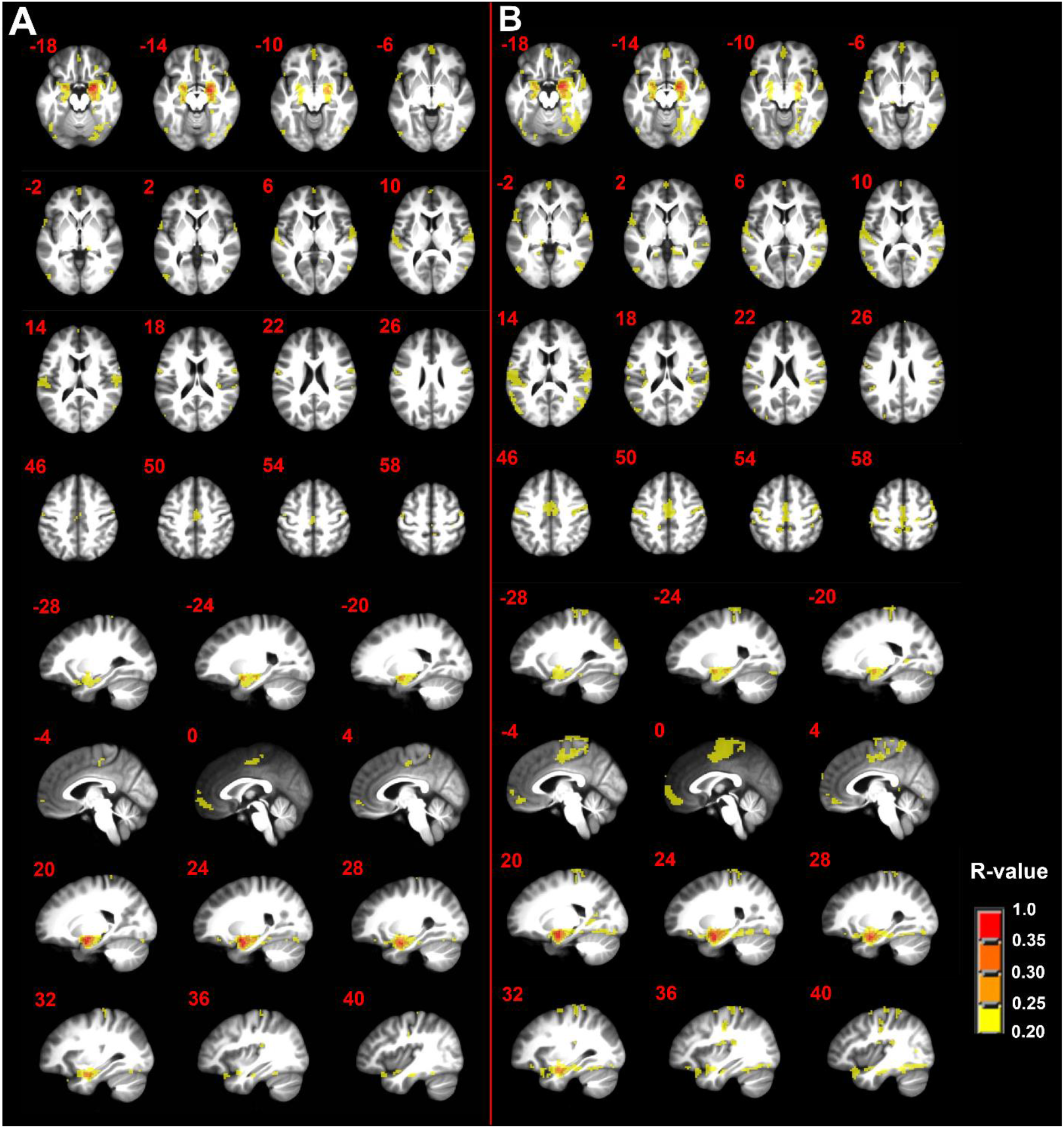
**A)** Seed-driven functional connectivity from right amygdala complex during the pain-free state and **B)** during the tonic pain state. n=50; minimum cluster size 108 mm^3^; p-value threshold 0.0001; R-value threshold 0.20. Axial and sagittal labels are in MNI coordinates.

### Supplemental Tables

**Supplemental Table 1.** Voxel table of seed driven functional connectivity network from left parabrachial complex during the pain-free state. Volume is in mm^3^ and maximum intensity is R value. Brain maps were threshold filtered at R=0.10.

| Cluster Number and Brain Region | Brodmann Area | Volume | Maximum Intensity | MNI Coordinates |
| --- | --- | --- | --- | --- |
| 1. Left Parabrachial Nucleus* |  | 38205 | 1.00 | (-11, -35, -32) |
| 2. Right Precuneus | 7 | 4860 | 0.120 | (2, -65, 56) |
| 3. Left Parieto-Occipital Sulcus | 7/19 | 3213 | 0.123 | (-5, -83, 44) |
| 4. Left Anterior Midcingulate Cortex | 32 | 2835 | 0.123 | (-2, 14, 41) |
| 5. Left Mediodorsal Thalamus |  | 1026 | 0.120 | (-5, -11, 11) |
| 6. Right Lingual Gyrus | 18 | 864 | 0.128 | (2, -92, -8) |
| 7. Left Midcingulate Cortex | 24 | 621 | 0.116 | (-2, -8, 44) |
| 8. Right Ventrolateral Thalamus |  | 540 | 0.119 | (11, -14, 8) |
| 9. Right Caudate |  | 540 | 0.119 | (11, 5, 8) |
| 10. Left Dorsal Superior Frontal Gyrus | 6 | 513 | 0.112 | (-2, 17, 62) |
| 11. Right Middle Frontal Gyrus | 9 | 324 | 0.109 | (56, 17, 35) |
| 12. Right Lingual Gyrus | 18 | 270 | 0.123 | (2, -80, -5) |
| 13. Right Cerebellar Declive |  | 216 | 0.107 | (17, -83, -23) |
| 14. Right Middle Occipital Gyrus | 18 | 216 | 0.110 | (29, -83, -17) |
| 15. Right Retrosplenial Cortex | 30 | 216 | 0.115 | (5, -44, 5) |
| 16. Right Paracentral Lobule | 5 | 216 | 0.106 | (14, -41, 53) |
| 17. Right Caudate Body |  | 189 | 0.127 | (17, -14, 20) |
| 18. Left Ventral Striatum |  | 162 | 0.123 | (-23, 5, -14) |
| 19. Left Superior Parietal Lobule | 7 | 162 | 0.114 | (-29, -80, 47) |
| 20. Left Inferior Parietal Lobule | 39 | 135 | 0.109 | (-32, -77, 35) |
| 21. Left Middle Cingulate Cortex | 23 | 108 | 0.116 | (-2, -17, 29) |
| 22. Left Postcentral Gyrus | 5 | 108 | 0.118 | (-8, -50, 77) |
| 23. Left Supplemental Motor Area | 6 | 108 | 0.107 | (-5, -8, 74) |

\* = includes seed region.

**Supplemental Table 2.** Voxel table of seed driven functional connectivity network from left parabrachial complex during the tonic pain state. Volume is in mm^3^ and maximum intensity is R value. Brain maps were threshold filtered at R=0.10.

| Cluster Number and Brain Region | Brodmann Area | Volume | Maximum Intensity | MNI Coordinates |
| --- | --- | --- | --- | --- |
| 1. Left Parabrachial Nucleus* |  | 126333 | 1.00 | (-11, -35, -32) |
| 2. Right Mediodorsal Thalamus |  | 8019 | 0.148 | (8, -14, 11) |
| 3. Right Precentral Gyrus | 6 | 3753 | 0.149 | (41, -14, 44) |
| 4. Left Inferior Frontal Operculum | 44 | 2808 | 0.136 | (-59, 8, 2) |
| 5. Right Superior Temporal Gyrus | 22 | 2349 | 0.122 | (65, -47, 20) |
| 6. Right Inferior Frontal Gyrus pars triangularis | 45 | 1458 | 0.129 | (56, 20, -5) |
| 7. Right Postcentral Gyrus | 40 | 1431 | 0.125 | (65, -20, 17) |
| 8. Left Postcentral Gyrus | 1/2/3 | 1323 | 0.135 | (-38, -20, 41) |
| 9. Left Middle Frontal Gyrus | 10 | 1134 | 0.126 | (-29, 50, 26) |
| 10. Left Central Sulcus | 4 | 1134 | 0.126 | (-23, -29, 65) |
| 11. Left Superior Occipital Gyrus | 19 | 810 | 0.116 | (-29, -83, 35) |
| 12. Left Superior Parietal Lobule | 7 | 810 | 0.119 | (29, -77, 50) |
| 13. Right Superior Frontal Gyrus | 10 | 756 | 0.117 | (29, 59, 5) |
| 14. Right Postcentral Gyrus | 1/2/3 | 729 | 0.119 | (20, -29, 65) |
| 15. Right Retrosplenial Cortex | 30 | 567 | 0.123 | (2, -44, 2) |
| 16. Left Medial Frontal Gyrus | 9 | 567 | 0.123 | (-2, 41, 29) |
| 17. Left Superior Temporal Gyrus | 22 | 513 | 0.116 | (-56, -59, 17) |
| 18. Right Middle Frontal Gyrus | 10 | 486 | 0.116 | (32, 50, 26) |
| 19. Right Precentral Gyrus | 6 | 459 | 0.116 | (62, 5, 11) |
| 20. Right Postcentral Gyrus | 1/2/3 | 351 | 0.110 | (23, -35, 77) |
| 21. Right Posterior Cingulate Gyrus | 31 | 297 | 0.121 | (5, -38, 26) |
| 22. Left Precuneus | 31 | 297 | 0.110 | (-14, -47, 35) |
| 23. Right Middle Frontal Gyrus | 9 | 297 | 0.112 | (53, 14, 38) |
| 24. Right Inferior Parietal Lobule | 40 | 297 | 0.109 | (44, -53, 53) |
| 25. Left Ventral Anterior Insula | 13 | 243 | 0.140 | (-38, 8, -14) |
| 26. Right Lateral Periaqueductal Gray/Superior Colliculus |  | 243 | 0.123 | (5, -23, -2) |
| 27. Left Parietal Operculum | 40 | 243 | 0.117 | (-48, -26, 14) |
| 28. Left Transverse Temporal Gyrus | 41/42 | 243 | 0.115 | (-62, -20, 14) |
| 29. Left Precentral Gyrus | 4 | 243 | 0.108 | (-56, -11, 47) |
| 30. Left Lingual Gyrus | 19 | 216 | 0.114 | (-17, -47, -2) |
| 31. Right Medial Frontal Gyrus | 10 | 216 | 0.114 | (2, 53, 5) |
| 32. Left Inferior Parietal Lobule | 40 | 216 | 0.106 | (-62, -41, 26) |
| 33. Right Superior Frontal Gyrus | 9 | 216 | 0.113 | (41, 35, 35) |
| 34. Left Superior Parietal Lobule | 7 | 216 | 0.107 | (-29, -65, 62) |
| 35. Right Superior Parietal Lobule | 7 | 216 | 0.110 | (20, -59, 71) |
| 36. Left Ventral Striatum |  | 189 | 0.122 | (-20, 5, -14) |
| **37. Left Lateral Periaqueductal Gray/Superior Colliculus** |  | **189** | **0.125** | **(-5, -20, -5)** |
| **38. Right Middle Temporal Gyrus** | **37** | **189** | **0.114** | **(59, -68, 5)** |
| **39. Left Middle Temporal Gyrus** | **37** | **189** | **0.111** | **(-56, -71, 11)** |
| **40. Left Inferior Semi-Lunar Lobule** |  | **162** | **0.111** | **(-44, -74, -44)** |
| **41. Right Ventral Striatum** |  | **162** | **0.117** | **(20, 5, -11)** |
| **42. Right Dorsal Posterior Insula** | **13** | **162** | **0.124** | **(38, -14, 20)** |
| **43. Left Middle Cingulate Cortex** | **24** | **135** | **0.115** | **(-11, -17, 41)** |
| **44. Right Parahippocampal Gyrus** | **35/36** | **108** | **0.116** | **(17, -35, -14)** |
| **45. Right Ventral Posterior Insula** | **13** | **108** | **0.114** | **(38, -14, -5)** |
| **46. Right Presubiculum** | **27** | **108** | **0.106** | **(17, -29, -5)** |
| **47. Left Ventral Middle Insula** | **13** | **108** | **0.113** | **(-41, -2, -2)** |
| **48. Left Retrosplenial Cortex** | **29** | **108** | **0.105** | **(-11, -44, 5)** |
| **49. Left Putamen** |  | **108** | **0.103** | **(-29, -23, 5)** |
| **50. Left Middle Occipital Gyrus** | **18** | **108** | **0.113** | **(-32, -98, 8)** |
| **51. Left Dorsal Posterior Insula** | **13** | **108** | **0.109** | **(-35, -32, 20)** |
| **52. Left Dorsal Posterior Insula** | **13** | **108** | **0.112** | **(-38, -17, 20)** |
| **53. Right Inferior Parietal Lobule** | **40** | **108** | **0.111** | **(50, -47, 23)** |
| **54. Right Middle Frontal Gyrus** | **9** | **108** | **0.106** | **(56, 20, 29)** |
| **55. Right Posterior Cingulate Cortex** | **31** | **108** | **0.107** | **(11, -41, 35)** |
| **56. Right Superior Frontal Gyrus** | **6** | **108** | **0.110** | **(26, -11, 74)** |

\* = includes seed region.

**Supplemental Table 3.** Voxel table of seed driven functional connectivity network from right parabrachial complex during the pain-free state. Volume is in mm^3^ and maximum intensity is R value. Brain maps were threshold filtered at R=0.10.

| Cluster Number and Brain Region | Brodmann Area | Volume | Maximum Intensity | MNI Coordinates |
| --- | --- | --- | --- | --- |
| **1. Right Parabrachial Nucleus\*** |  | **196263** | **1.00** | **(11, -35, -29)** |
| Left Parabrachial Nucleus |  |  | 0.265 | (-14, -35, -29) |
| Midline Basilar Pons |  |  | 0.168 | (-2, -20, -23) |
| Right Cuneus | 19 |  | 0.165 | (2, -83, 41) |
| Left Fusiform Gyrus | 37 |  | 0.161 | (-29, -74, -17) |
| **2. Right Caudate** |  | **11043** | **0.152** | **(8, 2, 8)** |
| **3. Left Precentral Gyrus** | **44** | **3564** | **0.140** | **(-62, 5, 5)** |
| **4. Right Inferior Frontal Gyrus (orbitalis)** | **47** | **1809** | **0.134** | **(53, 23, -5)** |
| **5. Left Precentral Gyrus** | **4** | **1620** | **0.132** | **(-38, -20, 47)** |
| **6. Right Middle Frontal Gyrus** | **46** | **1512** | **0.132** | **(53, 20, 32)** |
| **7. Right Lingual Gyrus** | **18** | **945** | **0.126** | **(23, -101, -11)** |
| **8. Left Transverse Temporal Gyrus** | **41/42** | **891** | **0.122** | **(-47, -29, 8)** |
| **9. Right Transverse Temporal Gyrus** | **41/42** | **837** | **0.126** | **(59, -14, 11)** |
| **10. Left Inferior Parietal Lobule** | **40** | **783** | **0.117** | **(-50, -41, 59)** |
| **11. Right Ventral Striatum** |  | **675** | **0.131** | **(26, 2, -11)** |
| **12. Right Precentral Gyrus** | **6** | **675** | **0.119** | **(41, -14, 38)** |
| **13. Left Middle Frontal Gyrus** | **6** | **648** | **0.142** | **(-38, -2, 65)** |
| **14. Left Superior Temporal Gyrus** | **22** | **540** | **0.112** | **(-65, -50, 8)** |
| **15. Right Temporoparietal Junction** | **22/40** | **513** | **0.117** | **(59, -50, 20)** |
| **16. Right Middle Frontal Gyrus** | **9** | **513** | **0.125** | **(-53, 20, 32)** |
| **17. Left Ventral Striatum** |  | **459** | **0.133** | **(-20, 5, -14)** |
| **18. Right Middle Frontal Gyrus** | **46** | **459** | **0.114** | **(53, 32, 23)** |
| **19. Right Supramarginal Gyrus** | **40** | **459** | **0.114** | **(53, -44, 38)** |
| **20. Right Postcentral Gyrus** | **1/2/3** | **432** | **0.123** | **(29, -38, 74)** |
| **21. Left Postcentral Gyrus** | **1/2/3** | **378** | **0.128** | **(-29, -38, 74)** |
| **22. Left Cerebellar Tonsil** |  | **351** | **0.137** | **(-26, -38, -44)** |
| **23. Left Middle Temporal Gyrus** | **21** | **324** | **0.124** | **(-68, -29, -2)** |
| **24. Right Dorsal Posterior Insula** | **13** | **324** | **0.128** | **(35, -29, 20)** |
| **25. Left Precentral Gyrus** | **4** | **324** | **0.113** | **(-59, -8, 23)** |
| **26. Right Cerebellar Tonsil** |  | **297** | **0.123** | **(26, -38, -44)** |
| **27. Left Inferior Semi-Lunar Lobule** |  | **297** | **0.115** | **(-41, -74, -44)** |
| **28. Right Middle Occipital Gyrus** | **19** | **297** | **0.120** | **(56, -71, 2)** |
| **29. Left Inferior Parietal Lobule** | **40** | **270** | **0.116** | **(-62, -41, 23)** |
| **30. Left Ventral Anterior Insula** | **13** | **243** | **0.138** | **(-35, 11, -17)** |
| **31. Right Superior Frontal Gyrus** | **10** | **243** | **0.107** | **(29, 68, 5)** |
| **32. Left Postcentral Gyrus** | **5** | **243** | **0.106** | **(-26, -50, 68)** |
| **33. Left Pulvinar Thalamus** |  | **216** | **0.126** | **(-14, -29, -2)** |
| **34. Left Ventral Posterior Insula** |  | **216** | **0.117** | **(-38, -20, -2)** |
| 35. Left Putamen |  | 216 | 0.113 | (-29, -20, 5) |
| 36. Left Middle Frontal Gyrus | 10 | 216 | 0.117 | (-29, 47, 26) |
| 37. Right Superior Frontal Gyrus | 9 | 216 | 0.114 | (44, 35, 35) |
| 38. Right Cerebellar Declive |  | 189 | 0.118 | (35, -83, -29) |
| 39. Right Substantia Nigra (pars reticulata) |  | 189 | 0.131 | (14, -26, -17) |
| 40. Right Angular Gyrus | 39 | 189 | 0.111 | (53, -56, 32) |
| 41. Right Superior Frontal Gyrus | 8 | 189 | 0.120 | (5, 32, 62) |
| 42. Right Putamen |  | 162 | 0.113 | (32, -14, 2) |
| 43. Left Middle Temporal Gyrus | 21/37 | 162 | 0.109 | (-53, -74, 14) |
| 44. Right Angular Gyrus | 39 | 162 | 0.109 | (47, -68, 35) |
| 45. Left Central Sulcus | 1/2/4 | 162 | 0.110 | (-23, -29, 71) |
| 46. Right Precentral Gyrus | 4 | 162 | 0.119 | (23, -26, 77) |
| 47. Left Lateral Geniculate Nucleus |  | 135 | 0.126 | (-23, -20, -8) |
| 48. Right Lateral Occipital Gyrus | 37 | 135 | 0.114 | (44, -74, 17) |
| 49. Right Cerebellar Tonsil |  | 108 | 0.107 | (17, -44, -44) |
| 50. Left Pyramis |  | 108 | 0.105 | (-35, -80, -41) |
| 51. Left Cerebellar Tonsil |  | 108 | 0.120 | (-53, -56, -35) |
| 52. Right Superior Semilunar Lobule |  | 108 | 0.104 | (32, -5, -32) |
| 53. Left Cerebellar Culmen |  | 108 | 0.119 | (-2, -59, -2) |
| 54. Right Periventricular Gray |  | 108 | 0.115 | (5, -23, -2) |
| 55. Right Superior Temporal Gyrus | 41/42 | 108 | 0.113 | (56, -29, 17) |
| 56. Left Temporoparietal Junction | 39 | 108 | 0.106 | (-53, -62, 23) |
| 57. Right Parieto-Occipital Fissure |  | 108 | 0.107 | (23, -59, 17) |
| 58. Right Dorsal Posterior Insula | 13 | 108 | 0.104 | (44, -32, 20) |
| 59. Right Middle Frontal Gyrus | 9 | 108 | 0.103 | (32, 56, 20) |
| 60. Left Middle Frontal Gyrus | 9 | 108 | 0.105 | (-32, 50, 32) |
| 61. Right Precentral Gyrus | 6 | 108 | 0.110 | (59, -8, 47) |
| 62. Left Precentral Gyrus | 6 | 108 | 0.115 | (-41, -5, 50) |
| 63. Left Precentral Gyrus | 4 | 108 | 0.118 | (-26, -29, 62) |
| 64. Right Precentral Gyrus | 4 | 108 | 0.116 | (20, -29, 65) |
| 65. Left Superior Frontal Gyrus | 8 | 108 | 0.117 | (-2, 26, 65) |

\* = includes seed region.

**Supplemental Table 4.** Voxel table of seed driven functional connectivity network from right parabrachial complex during the tonic pain state. Volume is in mm^3^ and maximum intensity is R value. Brain maps were threshold filtered at R=0.10.

| Cluster Number and Brain Region | Brodmann Area | Volume | Maximum Intensity | MNI Coordinates |
| --- | --- | --- | --- | --- |
| **1. Right Parabrachial Nucleus\*** |  | **470313** | **1.00** | **(11, -35, -29)** |
| Left Parabrachial Nucleus |  |  | 0.260 | (-14, -35, -29) |
| Left Rostral Medial Pons |  |  | 0.194 | (-2, -20, -23) |
| Left Anterior Middle Cingulate Cortex | 32 |  | 0.185 | (-2, 14, 41) |
| Right Medial Postcentral Gyrus | 5 |  | 0.183 | (2, -47, 68) |
| Right Cuneus | 17 |  | 0.180 | (2, -89, 5) |
| Left Inferior Occipital Gyrus | 19 |  | 0.179 | (-26, -86, -20) |
| Left Inferior Frontal Gyrus (orbitalis) | 45 |  | 0.177 | (-53, 20, -8) |
| Left Lingual Gyrus | 18 |  | 0.176 | (-2, -92, -8) |
| Right Postcentral Gyrus | 40 |  | 0.176 | (65, -23, 17) |
| Right Inferior Frontal Gyrus | 45 |  | 0.174 | (56, 20, -5) |
| Right Lingual Gyrus | 18 |  | 0.172 | (2, -80, -5) |
| Right Paracentral Lobule | 4 |  | 0.172 | (2, -17, 47) |
| Right Caudate |  |  | 0.169 | (14, 2, 14) |
| Left Cuneus | 17 |  | 0.167 | (-8, -104, -5) |
| Right Cerebellar Culmen |  |  | 0.165 | (17, -65, -14) |
| Left Parietal Operculum | 40 |  | 0.165 | (-62, -20, 14) |
| Left Postcentral Gyrus | 1/2/3 |  | 0.164 | (-8, -47, 77) |
| Right Mediodorsal Thalamus |  |  | 0.161 | (8, -17, 11) |
| Left Calcarine Gyrus | 17 |  | 0.159 | (-5, -68, 11) |
| Left Mediodorsal Thalamus |  |  | 0.158 | (-8, -17, 8) |
| **2. Left Superior Frontal Gyrus** | **10** | **6561** | **0.154** | **(-32, 53, 29)** |
| **3. Left Inferior Semilunar Lobule** |  | **1323** | **0.120** | **(-44, -74, -44)** |
| **4. Left Middle Frontal Gyrus** | **9** | **810** | **0.123** | **(-56, 14, 32)** |
| **5. Left Middle Frontal Gyrus** | **9** | **351** | **0.122** | **(-47, 26, 41)** |
| **6. Right Orbitofrontal Gyrus** | **47** | **297** | **0.127** | **(35, 38, -17)** |
| **7. Left Postcentral Gyrus** | **1/2/3** | **270** | **0.118** | **(-65, -20, 38)** |
| **8. Left Hippocampus** |  | **243** | **0.113** | **(-32, -23, -20)** |
| **9. Right Inferior Semilunar Lobule** |  | **216** | **0.110** | **(44, -68, -44)** |
| **10. Right Ventral Posterior Insula** |  | **216** | **0.118** | **(41, -2, -17)** |
| **11. Right Middle Temporal Gyrus** | **38** | **189** | **0.112** | **(44, 11, -47)** |
| **12. Right Superior Frontal Gyrus** | **11** | **189** | **0.115** | **(23, 47, -17)** |
| **13. Hypothalamus** |  | **189** | **0.112** | **(-5, -11, -11)** |
| **14. Right Cerebellar Tonsil** |  | **162** | **0.111** | **(11, -47, -44)** |
| **15. Left Hippocampus** |  | **162** | **0.107** | **(-26, -11, -23)** |
| **16. Left Superior Frontal Gyrus** | **11** | **162** | **0.111** | **(-26, 47, -17)** |
| **17. Left Superior Temporal Gyrus** | **21** | **162** | **0.112** | **(-68, -23, -2)** |
| **18. Left Superior Frontal Gyrus** | **10** | **162** | **0.123** | **(-17, 71, 17)** |
| 19. Right Superior Frontal Gyrus | 8 | 108 | 0.108 | (23, 17, 68) |
| 20. Right Cerebellar Tonsil |  | 108 | 0.120 | (20, -41, -47) |
| 21. Right Inferior Temporal Gyrus | 20 | 108 | 0.106 | (47, -2, -35) |
| 22. Right Parahippocampal Gyrus | 28 | 108 | 0.109 | (35, -11, -29) |
| 23. Left Superior Orbital Gyrus | 11 | 108 | 0.107 | (-14, 35, -17) |
| 24. Left Middle Orbital Gyrus | 47 | 108 | 0.113 | (-44, 56, -8) |
| 25. Bilateral Inferior Colliculus |  | 108 | 0.108 | (-2, -38, -8) |
| 26. Right Cerebellar Culmen |  | 108 | 0.112 | (2, -56, -2) |
| 27. Right Superior Temporal Gyrus | 22 | 108 | 0.107 | (50, -29, -2) |
| 28. Left Dorsal Anterior Insula | 13 | 108 | 0.108 | (-35, 11, 8) |
| 29. Left Inferior Frontal Gyrus (opercularis) | 44 | 108 | 0.106 | (-62, 11, 14) |
| 30. Left Temporoparietal Junction | 39 | 108 | 0.113 | (-50, -62, 26) |

\* = includes seed region.

**Supplemental Table 5.**
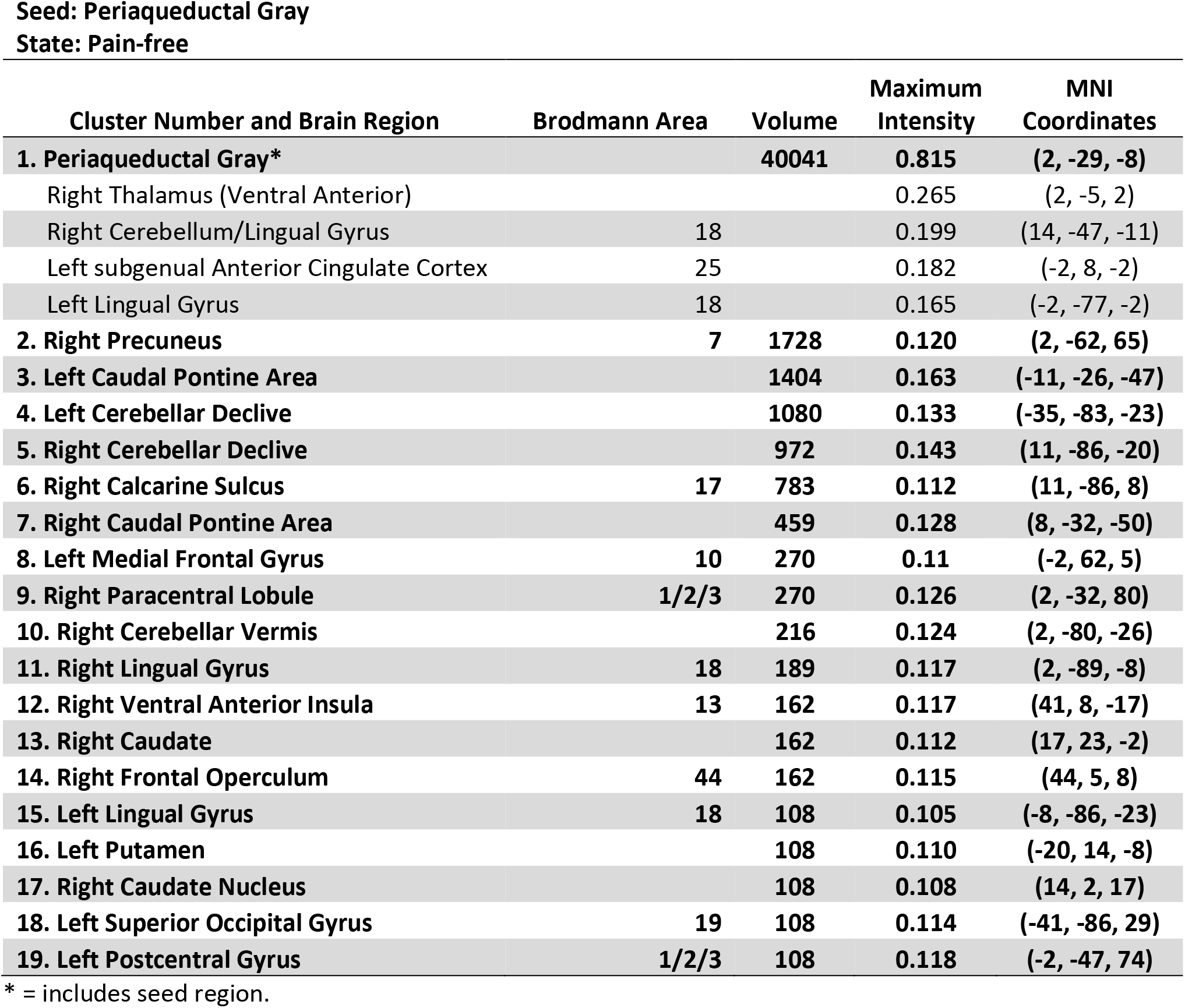
Voxel table of seed driven functional connectivity network from the periaqueductal gray area during the pain-free state. Volume is in mm^3^ and maximum intensity is R value. Brain maps were threshold filtered at R=0.10.

| Cluster Number and Brain Region | Brodmann Area | Volume | Maximum Intensity | MNI Coordinates |
| --- | --- | --- | --- | --- |
| **1. Periaqueductal Gray\*** |  | **40041** | **0.815** | **(2, -29, -8)** |
| Right Thalamus (Ventral Anterior) |  |  | 0.265 | (2, -5, 2) |
| Right Cerebellum/Lingual Gyrus | 18 |  | 0.199 | (14, -47, -11) |
| Left subgenual Anterior Cingulate Cortex | 25 |  | 0.182 | (-2, 8, -2) |
| Left Lingual Gyrus | 18 |  | 0.165 | (-2, -77, -2) |
| **2. Right Precuneus** | **7** | **1728** | **0.120** | **(2, -62, 65)** |
| **3. Left Caudal Pontine Area** |  | **1404** | **0.163** | **(-11, -26, -47)** |
| **4. Left Cerebellar Declive** |  | **1080** | **0.133** | **(-35, -83, -23)** |
| **5. Right Cerebellar Declive** |  | **972** | **0.143** | **(11, -86, -20)** |
| **6. Right Calcarine Sulcus** | **17** | **783** | **0.112** | **(11, -86, 8)** |
| **7. Right Caudal Pontine Area** |  | **459** | **0.128** | **(8, -32, -50)** |
| **8. Left Medial Frontal Gyrus** | **10** | **270** | **0.11** | **(-2, 62, 5)** |
| **9. Right Paracentral Lobule** | **1/2/3** | **270** | **0.126** | **(2, -32, 80)** |
| **10. Right Cerebellar Vermis** |  | **216** | **0.124** | **(2, -80, -26)** |
| **11. Right Lingual Gyrus** | **18** | **189** | **0.117** | **(2, -89, -8)** |
| **12. Right Ventral Anterior Insula** | **13** | **162** | **0.117** | **(41, 8, -17)** |
| **13. Right Caudate** |  | **162** | **0.112** | **(17, 23, -2)** |
| **14. Right Frontal Operculum** | **44** | **162** | **0.115** | **(44, 5, 8)** |
| **15. Left Lingual Gyrus** | **18** | **108** | **0.105** | **(-8, -86, -23)** |
| **16. Left Putamen** |  | **108** | **0.110** | **(-20, 14, -8)** |
| **17. Right Caudate Nucleus** |  | **108** | **0.108** | **(14, 2, 17)** |
| **18. Left Superior Occipital Gyrus** | **19** | **108** | **0.114** | **(-41, -86, 29)** |
| **19. Left Postcentral Gyrus** | **1/2/3** | **108** | **0.118** | **(-2, -47, 74)** |

\* = includes seed region.

**Supplemental Table 6.** Voxel table of seed driven functional connectivity network from the periaqueductal gray area during the tonic pain state. Volume is in mm^3^ and maximum intensity is R value. Brain maps were threshold filtered at R=0.10.

| Cluster Number and Brain Region | Brodmann Area | Volume | Maximum Intensity | MNI Coordinates |
| --- | --- | --- | --- | --- |
| **1. Periaqueductal Gray\*** |  | **46575** | **0.823** | **(2, -29, -8)** |
| Ventral Anterior Thalamus |  |  | 0.221 | (2, -5, 2) |
| **2. Left Anterior Midcingulate Cortex** | **32** | **5292** | **0.150** | **(-2, 26, 32)** |
| **3. Right Lingual Gyrus** | **18** | **4563** | **0.160** | **(2, -80, -5)** |
| **4. Right Cerebellar Declive** |  | **2106** | **0.127** | **(2, -80, -29)** |
| **5. Left Precuneus** | **7** | **1404** | **0.122** | **(-5, -62, 56)** |
| **6. Right Cerebellar Declive** |  | **918** | **0.131** | **(20, -83, -23)** |
| **7. Left Caudal Pontine Area** |  | **891** | **0.129** | **(-11, -26, -44)** |
| **8. Right Rostral Lateral Medulla (+)** |  | **540** | **0.140** | **(11, -35, -47)** |
| **9. Right Rostral Ventral Medulla (#)** |  | **459** | **0.141** | **(5, -23, -47)** |
| **10. Left Paracentral Lobule** |  | **432** | **0.119** | **(-8, -44, 53)** |
| **11. Left Paracentral Lobule (Crown)** | **1/2/3** | **405** | **0.130** | **(-2, -41, 77)** |
| **12. Cerebellar Vermis** |  | **351** | **0.121** | **(-2, -53, -35)** |
| **13. Left Postcentral Gyrus** | **5/7** | **351** | **0.123** | **(-2, -53, 74)** |
| **14. Left Lingual Gyrus** | **18** | **324** | **0.115** | **(-2, 68, 5)** |
| **15. Left Superior Occipital Gyrus** | **19** | **216** | **0.121** | **(-41, -86, 26)** |
| **16. Right Cuneus** | **19** | **216** | **0.124** | **(2, -89, 38)** |
| **17. Right Middle Cingulate Cortex** | **24** | **189** | **0.108** | **(2, -11, 38)** |
| **18. Right Mid Lateral Medulla (%)** |  | **108** | **0.116** | **(11, -41, -56)** |
| **19. Left Cerebellar Declive** |  | **108** | **0.110** | **(-35, -83, -26)** |
| **20. Left Cerebellar Declive** |  | **108** | **0.109** | **(-29, -77, -23)** |
| **21. Right Ventral Anterior Insula** | **13** | **108** | **0.118** | **(41, 5, -17)** |
| **22. Left Posterior Cingulate Cortex** | **29** | **108** | **0.113** | **(-8, -47, 8)** |
| **23. Left Ventral Lateral Thalamus** |  | **108** | **0.105** | **(-17, -11, 17)** |

\* = includes seed region. + = including facial nucleus, spinal trigeminal nucleus, cochlear nucleus, # including raphe nuclei, hypoglossal nucleus, dorsal motor nucleus, % including lateral reticular nucleus and spinal trigeminal nucleus

**Supplemental Table 7.** Voxel table of seed driven functional connectivity network from left amygdala complex during the pain-free state. Volume is in mm^3^ and maximum intensity is R value. Brain maps were threshold filtered at R=0.20.

| Cluster Number and Brain Region | Brodmann Area | Volume | Maximum Intensity | MNI Coordinates |
| --- | --- | --- | --- | --- |
| 1. Left Amygdala* |  | 16119 | 0.687 | (-20, -5, -17) |
| 2. Right Amygdala |  | 5994 | 0.366 | (17, -5, -17) |
| 3. Right Ventral Medial Frontal Cortex | 10 | 2997 | 0.267 | (2, 62, -5) |
| 4. Left Frontal Operculum | 43 | 2916 | 0.263 | (-62, 5, 2) |
| 5. Right Frontal Operculum | 43 | 2808 | 0.243 | (65, -2, 8) |
| 6. Left Paracentral Lobule | 4 | 2160 | 0.243 | (-2, -20, 53) |
| 7. Left Fusiform Gyrus | 37 | 837 | 0.245 | (-50, -65, -17) |
| 8. Right Postcentral Gyrus | 1/2/3 | 810 | 0.225 | (35, -29, 71) |
| 9. Left Precentral Gyrus | 4 | 486 | 0.234 | (-62, -5, 23) |
| 10. Right Dorsal Posterior Insula | 13 | 459 | 0.243 | (38, -29, 20) |
| 11. Right Precentral Gyrus | 6 | 432 | 0.232 | (62, -2, 23) |
| 12. Right Central Sulcus | 4 | 432 | 0.221 | (14, -32, 68) |
| 13. Right Paracentral Lobule | 4 | 405 | 0.221 | (2, -29, 74) |
| 14. Right Precentral Gyrus | 4 | 378 | 0.229 | (47, -14, 62) |
| 15. Right Paracentral Lobule | 5 | 324 | 0.219 | (5, -44, 62) |
| 16. Right Middle Temporal Gyrus | 21 | 243 | 0.225 | (59, -2, -14) |
| 17. Left Middle Temporal Gyrus | 21 | 162 | 0.224 | (-65, -11, -8) |
| 18. Left Middle Occipital Gyrus | 19 | 162 | 0.212 | (-53, -77, -2) |
| 19. Right Mediodorsal Thalamus |  | 162 | 0.214 | (2, -8, 8) |
| 20. Left Temporo-occipital Junction | 39 | 162 | 0.217 | (-56, -71, 11) |
| 21. Left Postcentral Gyrus | 1/2/3 | 135 | 0.228 | (-35, -38, 74) |
| 22. Right Fusiform Gyrus | 37 | 108 | 0.208 | (44, -56, -20) |
| 23. Left Parahippocampal Gyrus | 36 | 108 | 0.216 | (-23, -41, -14) |
| 24. Right Precentral Gyrus | 6 | 108 | 0.215 | (59, -8, 44) |

\* = includes seed region.

**Supplemental Table 8.** Voxel table of seed driven functional connectivity network from left amygdala complex during the tonic pain state. Volume is in mm^3^ and maximum intensity is R value. Brain maps were threshold filtered at R=0.20.

| Cluster Number and Brain Region | Brodmann Area | Volume | Maximum Intensity | MNI Coordinates |
| --- | --- | --- | --- | --- |
| **1. Left Amygdala\*** |  | **49599** | **0.689** | **(-20, -2, -20)** |
| Left Superior Temporal Gyrus | 22 |  | 0.298 | (-59, 11, -2) |
| **2. Left Middle Cingulate Cortex** | **24** | **12285** | **0.257** | **(-5, -11, 47)** |
| **3. Right Superior Temporal Gyrus** | **22** | **9747** | **0.299** | **(62, -2, 5)** |
| Right Parietal Operculum | 40 |  | 0.279 | (68, -20, 17) |
| Right Dorsal Posterior Insula | 13 |  | 0.276 | (41, -32, 20) |
| **4. Right Precentral Gyrus** | **6** | **8046** | **0.263** | **(46, -14, 62)** |
| **5. Right Middle Temporal Gyrus** | **37** | **6912** | **0.266** | **(56, -65, 8)** |
| **6. Right Amygdala** |  | **5616** | **0.349** | **(20, -2, -17)** |
| **7. Left Precentral Gyrus** | **4** | **5589** | **0.260** | **(-41, -20, 68)** |
| **8. Left Ventral Medial Frontal Cortex** | **10** | **4536** | **0.252** | **(-2, 56, -17)** |
| **9. Left Postcentral Gyrus** | **1/2/3** | **1863** | **0.251** | **(-35, -41, 71)** |
| **10. Right Superior Temporal Gyrus** | **22** | **1485** | **0.246** | **(62, -50, 17)** |
| **11. Right Retrosplenial Complex** | **29** | **1080** | **0.229** | **(8, -41, 2)** |
| **12. Right Superior Parietal Lobule** | **7** | **648** | **0.214** | **(17, -62, 68)** |
| **13. Left Lingual Gyrus** | **19** | **513** | **0.220** | **(-17, -50, 2)** |
| **14. Right Postcentral Gyrus** | **1/2/3** | **459** | **0.234** | **(56, -23, 56)** |
| **15. Left Superior Parietal Lobule** | **7** | **378** | **0.212** | **(-26, -59, 71)** |
| **16. Right Fusiform Gyrus** | **37** | **351** | **0.215** | **(20, -65, -14)** |
| **17. Right Superior Temporal Gyrus** | **22** | **351** | **0.242** | **(59, -5, -11)** |
| **18. Left Precuneus** | **19** | **351** | **0.222** | **(-29, -86, 32)** |
| **19. Right Inferior Frontal Gyrus (orbitalis)** | **47** | **297** | **0.214** | **(41, 26, -20)** |
| **20. Left Middle Frontal Gyrus** | **8** | **270** | **0.242** | **(-56, 8, 41)** |
| **21. Left Thalamus (Pulvinar)** |  | **243** | **0.221** | **(-14, -32, -2)** |
| **22. Right Parahippocampal Gyrus** | **36** | **216** | **0.247** | **(26, -32, -20)** |
| **23. Left Retrosplenial Complex** | **29** | **216** | **0.226** | **(-8, -44, 5)** |
| **24. Left Dorsal Posterior Insula** | **13** | **216** | **0.255** | **(-38, -14, 20)** |
| **25. Left Fusiform Gyrus** | **37** | **189** | **0.220** | **(-35, -41, -23)** |
| **26. Right Inferior Frontal Gyrus (orbitalis)** | **47** | **162** | **0.216** | **(29, 20, -23)** |
| **27. Right Posterior Insula** | **13** | **162** | **0.235** | **(38, -23, 5)** |
| **28. Right Superior Temporal Gyrus** | **22** | **162** | **0.208** | **(47, -41, 11)** |
| **29. Left Superior Parietal Lobule** | **7** | **162** | **0.214** | **(-26, -65, 65)** |
| **30. Right Superior Temporal Gyrus** | **38** | **135** | **0.230** | **(53, 17, -23)** |
| **31. Right Superior Temporal Gyrus** | **38** | **108** | **0.210** | **(59, 11, -17)** |
| **32. Right Inferior Frontal Gyrus (orbitalis)** | **47** | **108** | **0.223** | **(35, 38, -17)** |
| **33. Right Lingual Gyrus** | **18** | **108** | **0.205** | **(11, -80, -11)** |
| **34. Left Middle Insula** | **13** | **108** | **0.216** | **(-41, -5, 2)** |
| **35. Left Middle Insula** | **13** | **108** | **0.219** | **(-38, -8, 5)** |
| **36. Right Posterior Cingulate Gyrus** | **30** | **108** | **0.206** | **(17, -56, 8)** |
| **37. Left Dorsal Posterior Insula** | **13** | **108** | **0.218** | **(-35, -26, 11)** |
| **38. Left Posterior Cingulate Cortex** | **23** | **108** | **0.222** | **(-5, -53, 20)** |
| **39. Left Dorsal Posterior Insula** | **13** | **108** | **0.244** | **(-38, -17, 20)** |
| **40. Right Medial Frontal Gyrus** | **10** | **108** | **0.208** | **(2, 65, 20)** |
| **41. Left Precuneus** | **7** | **108** | **0.214** | **(-5, -56, 50)** |

\* = includes seed region.

**Supplemental Table 9.**
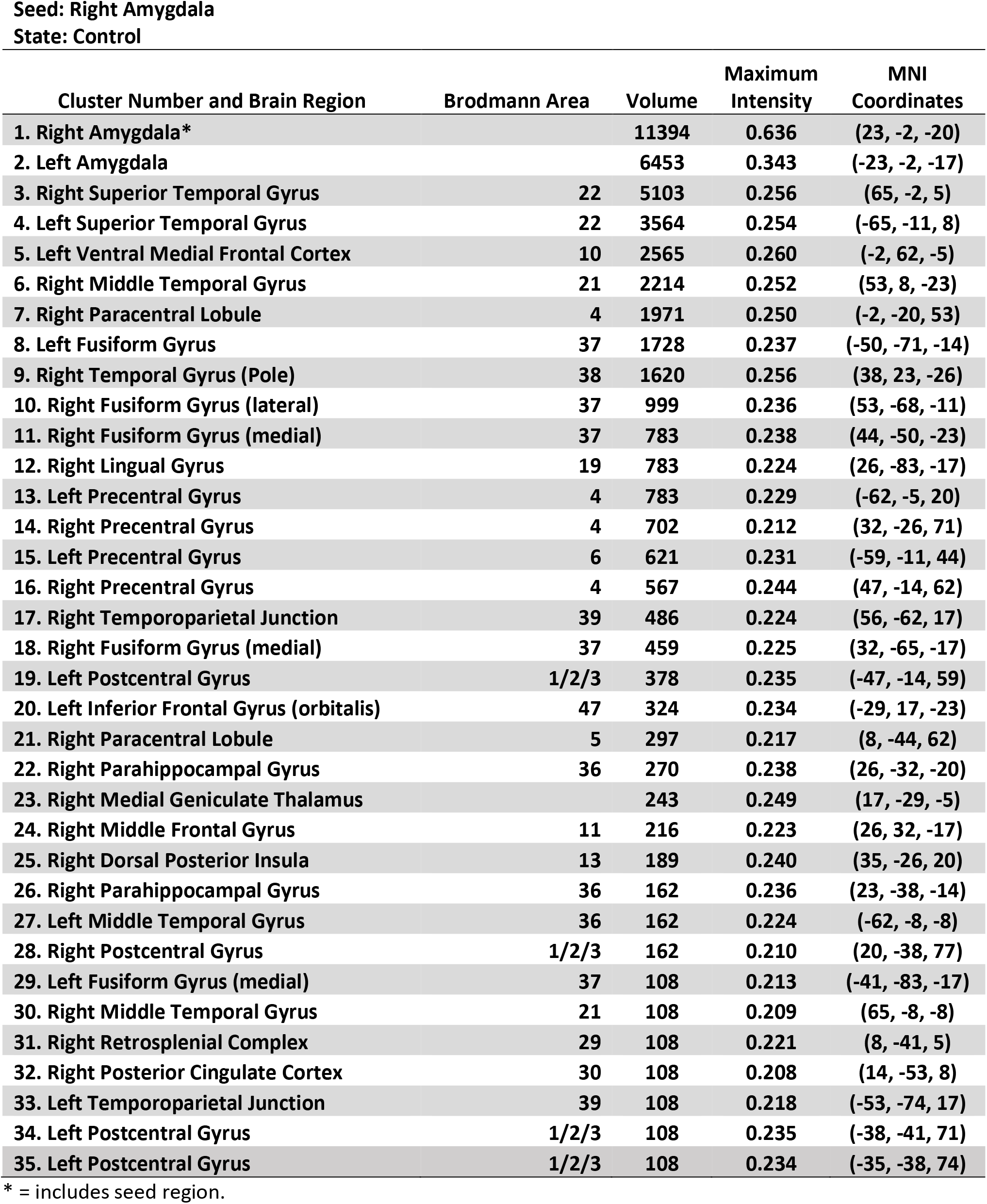
Voxel table of seed driven functional connectivity network from right amygdala complex during the pain-free state. Volume is in mm^3^ and maximum intensity is R value. Brain maps were threshold filtered at R=0.20.

**Supplemental Table 10.** Voxel table of seed driven functional connectivity network from right amygdala complex during the tonic pain state. Volume is in mm^3^ and maximum intensity is R value. Brain maps were threshold filtered at R=0.20.

| Cluster Number and Brain Region | Brodmann Area | Volume | Maximum Intensity | MNI Coordinates |
| --- | --- | --- | --- | --- |
| **1. Right Amygdala\*** |  | **44820** | **0.641** | **(23, -2, -20)** |
| Right Superior Temporal Gyrus | 22 |  | 0.303 | (62, 2, 5) |
| Right Superior Temporal Gyrus (Pole) | 38 |  | 0.292 | (41, 23, -23) |
| Right Middle Occipital Gyrus | 37 |  | 0.279 | (50, -68, -14) |
| Right Middle Temporal Gyrus | 37 |  | 0.270 | (53, -68, 5) |
| Right Inferior Frontal Gyrus (orbitalis) | 47 |  | 0.265 | (50, 20, -11) |
| Right Fusiform Gyrus | 37 |  | 0.265 | (47, -49, -20) |
| **2. Left Paracentral Lobule** | **4** | **30078** | **0.262** | **(-2, -23, 56)** |
| **3. Left Superior Temporal Gyrus** | **22** | **9801** | **0.281** | **(-59, 11, -2)** |
| Left Inferior Frontal Gyrus | 47 |  | 0.263 | (-50, 20, -11) |
| **4. Left Middle Occipital Gyrus** | **19** | **7020** | **0.258** | **(-52, -77, 2)** |
| **5. Left Amygdala** |  | **6264** | **0.338** | **(-26, 2, -14)** |
| **6. Left Ventral Medial Frontal Cortex** | **10** | **3537** | **0.273** | **(-2, 53, -17)** |
| **7. Right Parahippocampal Gyrus** | **36** | **1539** | **0.239** | **(20, -47, -2)** |
| **8. Right Dorsal Posterior Insula** | **13** | **918** | **0.262** | **(41, -32, 20)** |
| **9. Right Superior Temporal Gyrus** | **22** | **810** | **0.234** | **(62, -50, 17)** |
| **10. Left Cuneus** | **19** | **513** | **0.219** | **(-29, -89, 29)** |
| **11. Right Middle Frontal Gyrus** | **47** | **405** | **0.233** | **(35, 38, -17)** |
| **12. Right Superior Temporal Gyrus** | **22** | **405** | **0.226** | **(47, -38, 5)** |
| **13. Left Parahippocampal Gyrus** | **36** | **378** | **0.233** | **(-17, -50, 2)** |
| **14. Right Middle Temporal Gyrus** | **22** | **378** | **0.219** | **(65, -35, 2)** |
| **15. Left Superior Temporal Gyrus (Pole)** | **38** | **324** | **0.230** | **(-50, 14, -29)** |
| **16. Right Dorsal Posterior Insula** | **13** | **270** | **0.245** | **(38, -11, 20)** |
| **17. Left Inferior Parietal Lobule** | **40** | **270** | **0.219** | **(-53, -35, 59)** |
| **18. Left Parahippocampal Gyrus** | **36** | **216** | **0.228** | **(-26, -35, -20)** |
| **19. Left Posterior Insula** | **13** | **216** | **0.240** | **(-41, -20, -2)** |
| **20. Left Dorsal Middle Insula** | **13** | **216** | **0.243** | **(-38, -17, 20)** |
| **21. Left Middle Frontal Gyrus** | **6** | **216** | **0.233** | **(-38, -5, 65)** |
| **22. Left Medial Geniculate Thalamus** |  | **189** | **0.231** | **(-14, -32, -2)** |
| **23. Left Superior Temporal Gyrus (Pole)** | **38** | **162** | **0.222** | **(-38, 20, -26)** |
| **24. Right Postcentral Gyrus** | **1/2/3** | **162** | **0.234** | **(59, -20, 50)** |
| **25. Left Lingual Gyrus** | **18** | **108** | **0.208** | **(-11, -80, -11)** |
| **26. Left Middle Temporal Gyrus** | **21** | **108** | **0.209** | **(-65, -14, -5)** |
| **27. Right Medial Frontal Gyrus** | **10** | **108** | **0.214** | **(5, 68, 23)** |

\* = includes seed region.

## Notes

### Competing Interest Statement

The authors have declared no competing interest.

